# Sporophyte Directed Gametogenesis via the Ubiquitin Proteasome System

**DOI:** 10.1101/2024.12.23.630054

**Authors:** Prakash Sivakumar, Saurabh Pandey, A Ramesha, Jayeshkumar Narsibhai Davda, Aparna Singh, Chandan Kumar, Hardik Gala, Veeraputhiran Subbiah, Harikrishna Adicherla, Jyotsna Dhawan, L. Aravind, Imran Siddiqi

## Abstract

Plants alternate between diploid sporophyte and haploid gametophyte generations. In mosses which retain features of ancestral land plants, the gametophyte is dominant and has an independent existence. However, in flowering plants the gametophyte has undergone evolutionary reduction to just a few cells enclosed within the sporophyte. The gametophyte is thought to retain genetic control of its development even after reduction. Here we demonstrate that male gametophyte development in Arabidopsis, long considered to be autonomous, is also under genetic control of the sporophyte via a repressive mechanism involving large-scale regulation of protein turnover. We identify an Arabidopsis gene *SHUKR* as an inhibitor of male gametogenesis. SHUKR is unrelated to proteins of known function and acts sporophytically in meiosis to control gametophyte development by negatively regulating expression of a large set of ubiquitination genes specific to post-meiotic gametogenesis. This control is late-emerging as *SHUKR* homologs are found only in eudicots. We show that *SHUKR* is rapidly evolving under positive selection suggesting that variation in control of protein turnover during male gametogenesis has played an important role in evolution within eudicots.

## Introduction

Alternation between two distinct generations: a diploid sporophyte and a haploid gametophyte is a central feature of the land plant life cycle^1–3^. Land plants likely evolved from an algal ancestor related to extant charophycean algae which comprise the closest relatives to land plants^4,5^. Charophycean algae have a predominantly haploid life cycle, the diploid phase being transient as the zygote directly undergoes meiosis to give haploid spores. The emergence of a multicellular diploid sporophyte contributed to reproductive success on land by increasing the number of spores produced from a single fertilization event, and also increased potential for genetic variability and evolutionary change^1^. From a simple ancestral body plan, land plants comprising bryophytes (mosses, liverworts, and hornworts) and vascular plants evolved progressively specialized cell types, organs, and architectures adapted for life on land. These acquisitions have been based on development and increase in complexity of a multicellular sporophyte^6,7^. In vascular plants the sporophyte evolved to become the dominant phase and the gametophyte has reduced. In flowering plants the gametophyte is reduced to just a few cells and develops enclosed within the sporophyte^8^ (Fig. 1a). The differing trajectories taken by the sporophyte and gametophyte in land plant evolution reflects an uncoupling between their body plan evolution consistent with different genetic programs directing their body organization^7,9,10,11,12^.

**Fig. 1:**
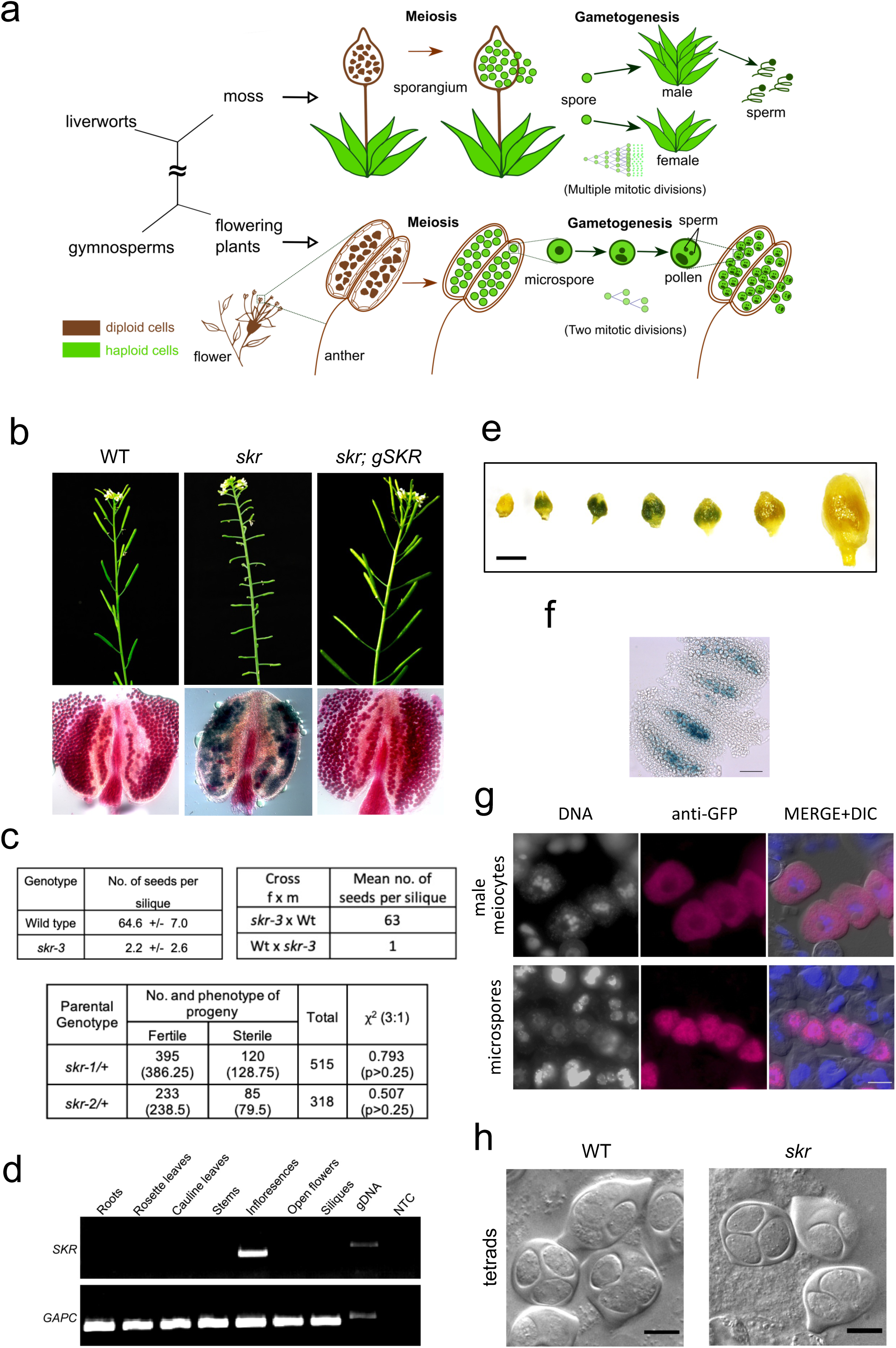
*SHUKR* is required for pollen development. (a) Schematic showing reduction of the gametophyte in flowering plants. (b) Sterility (short siliques) and reduced pollen viability (green staining) in *skr*, and complementation by a genomic *SKR* transgene. (c) Tables showing reduced seed set, male-specific sterility, and recessive Mendelian segregation for *skr*; numbers in brackets are those expected for a 3:1 ratio and the χ^2^ test p value. (d) RT-PCR analysis of SKR expression in vegetative and reproductive tissues. (e) GUS staining for *proSKR:GFP-GUS* transgene expression in successive unopened buds. (f) GUS staining for *proSKR:GFP-GUS* expression in meiotic stage anthers. (g) anti-GFP antibody staining for *proSKR:GFP-GUS* expression in male meiosis and microspores. (h) Normal tetrad formation in *skr*. Scale bar: (e) 0.5 mm, (f) 20 µm.

Formation of a germline is an early step in animal development^13,14^, however in plants the formation of germ cell precursors takes place late in development^15^. The fact that bryophytes have a multicellular independent gametophyte which produces gametes following extensive mitotic divisions allows overall assignment of the germline to the gametophyte generation^16^. In flowering plants the assignment of the germline is more difficult because of extreme reduction of the gametophyte and its development enclosed within the sporophyte^15,17,18^. Evidence from comparative evolution showing functional conservation of transcription factors controlling early steps in germ cell development between the liverwort *Marchantia polymorpha* and Arabidopsis argues for the male germline residing in the gametophyte^16^. Asymmetric cell division of the haploid microspore in Arabidopsis at pollen mitosis I forms a generative cell (GC) and vegetative cell (VC)^16,18^. In Arabidopsis, specification of the GC requires bHLH transcription factors, BNB1/BNB2 and DROP1/DROP2 which act in the gametophyte and are orthologous to MpBNB and MpDROP respectively, that also act in the gametophyte to direct gametangia specification and differentiation in Marchantia^19,20^. Sperm differentiation is controlled by gametophytic action of the bHLH transcription factor DUO1 in both Arabidopsis and Marchantia^21,22^. On the female side gametophytically acting transcription, splicing, and signaling related factors are important for cell specification and cellularization in Arabidopsis female gametophytes^23–29^. Therefore, the gametophyte in flowering plants has been considered to largely retain genetic control of its development even though it lacks an independent existence.

The extreme reduction of the gametophyte and its development enclosed within the sporophyte provided a “functionally unitary individual” for natural selection^30^, addressing the sensitive requirements for fertilization and reproductive development in a land environment. Formation and function of meiotic cells depends on a range of cellular interactions in the sporophyte^8,31–33^. These interactions involve transcriptional regulators acting nonautonomously^34–36^, cell surface receptor signaling^37–39^, and small RNA pathways^36,40,41^ together with phytohormone effectors^42^. Small RNAs^43^ and the phytohormone auxin^44,45^ have also been shown to mediate developmental interactions between cells within gametophytes. The tapetum is a sporophytic nurse tissue surrounding male meiocytes and provides enzymes for release of microspores and components for synthesis of the protective secondary outer coat of the pollen grain, the exine^46,47^. Mutants deficient in tapetal function manifest defects in exine formation and lead to pollen abortion^48–50^. The tapetum also produces small RNAs that direct DNA methylation of transposons and selected genes in male meiosis contributing to genome integrity and paternal epigenetic inheritance^51^.

Studies have shown that isolated uninucleate microspores can form functional pollen in vitro suggesting their development into pollen is autonomous^52,53^. It remains unclear whether genetic controls during meiosis broadly regulate postmeiotic gametophyte development. A challenge in uncovering such regulation has been the existence of very many sporophytic mutants that affect core meiosis functions including meiotic recombination and chromosome organization^54,55^ or else development of somatic cells within the anther or ovule^56–59^. These mutants have direct or indirect effects on gametophyte development and show spore or gametophyte abortion, making it difficult to distinguish from mutants affecting meiotic regulation of postmeiotic gametogenesis. Here we report the discovery in Arabidopsis that following evolutionary reduction in cell number of the gametophyte, the sporophyte within the eudicot lineage has gone on to evolve a process established in meiosis under control of the *SHUKR* gene, by which it directs male gametophyte development. The *shukr* mutant shows normal meiosis but postmeiotic development is defective. This process controlled by SHUKR involves large-scale control of regulated protein turnover in the developing gametophyte and is under rapid evolution with positive selection. Our findings suggest that the sporophyte in addition to providing supportive conditions for gametogenesis and fertilization, also exerts genetic control in meiosis of postmeiotic male gametogenesis. This control is part of a process that is capable of generating variation in the gametophyte during evolution, which has adaptive value.

## Results

### *SHUKR* is required for male fertility

We identified the *SHUKR* gene in a screen for novel meiotic functions based on coexpression with known meiotic genes in Arabidopsis (see Methods). Three independent insertion lines having insertions in *At2g38690* showed male sterility (Fig. 1b, Extended Data Fig. 1a). Allelism tests indicated the mutants to be allelic. Anthers from plants homozygous for the insertion contained a highly reduced number of viable pollen, resulting in sterility (Fig. 1b, c). Use of wild type pollen as the male donor in crosses to mutant plants resulted in full seed set indicating that female fertility was unaffected (Fig. 1c). The male sterile phenotype was inherited as a single gene recessive trait (Fig. 1c, Extended Data Fig. 1c). We named the *At2g38690* gene *SHUKR* (*SKR*) after the Sanskrit word Shukranu (/śukrāṇu/) for sperm. We confirmed the phenotype was due to loss of function of *SKR* by complementation with a transgenic copy. *skr/+* plants were transformed with wild type *At2g38690* genomic DNA (*gSKR*) by in planta transformation and T1 *skr* progeny were examined for fertility. The *gSKR* transgene rescued male sterility confirming the mutant phenotype was due to disruption of *At2g38690* (Fig. 1b, Extended Data Fig. 1d). Examination of SKR gene expression based on tissue RT-PCR, a proSKR:GFP-GUS reporter, and analysis of publicly available expression profiling data showed that SKR gene expression was specific to reproductive tissues and more specifically to male meiocytes (Fig. 1d-g; Extended Data Fig. 2). Expression of SKR protein was also meiosis-specific (Fig. 2c).

**Fig. 2:**
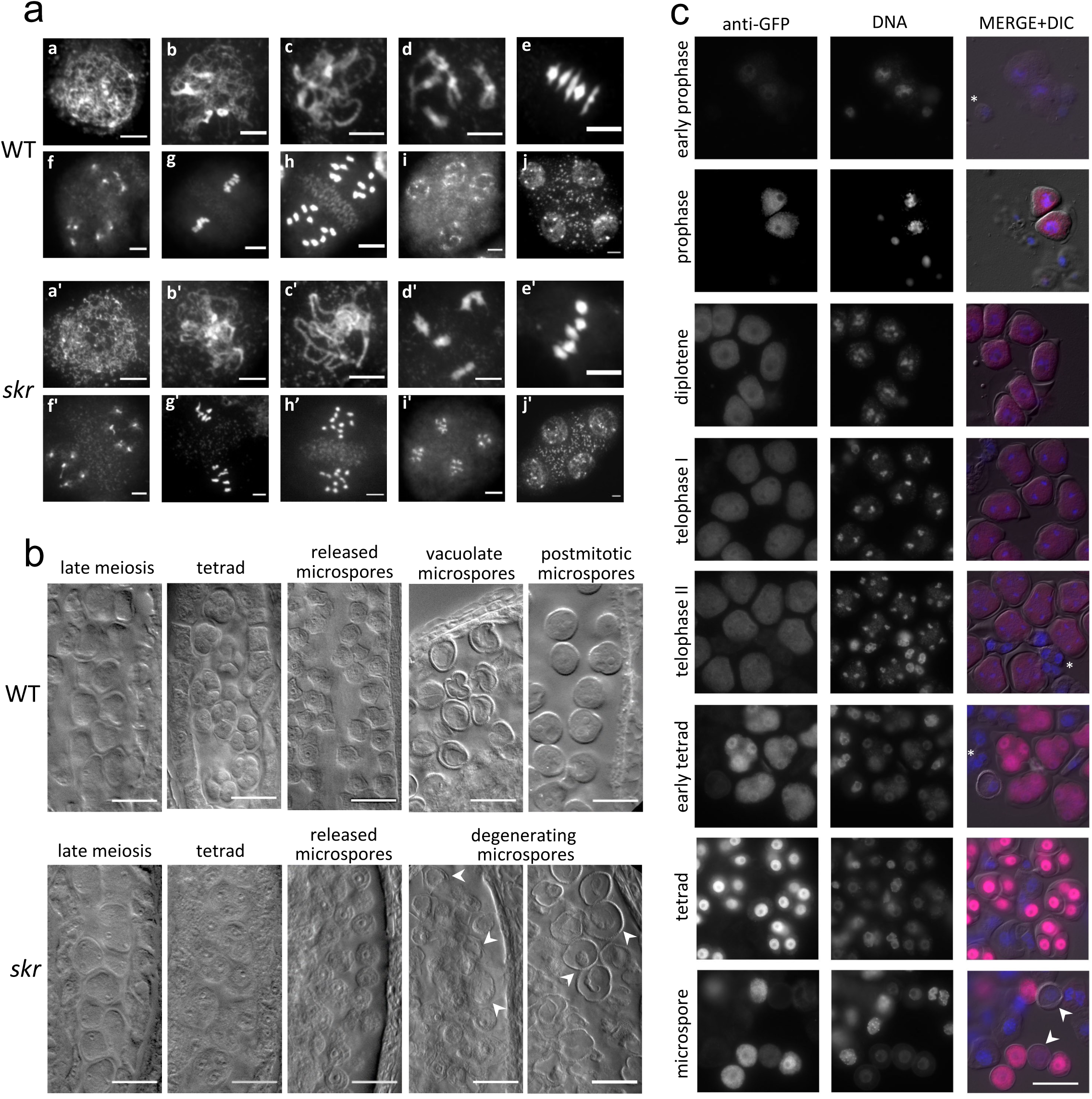
Staging of *skr* phenotype and SKR protein expression. (a) Meiotic chromosome development and segregation in wild type (a-j) and corresponding stages in *skr* (a’-j’). a-d,a’-d’ prophase I; e,e’ metaphase I; f,f’ telophase I; g,g’ metaphase II; h,h’ anaphase II; i,i’ telophase II; j,j’ tetrad. 5846 meiocytes were scored for *skr* and 2281 for WT. (b) Whole mount DIC images of cleared anthers from successive stage buds; arrowheads indicate degenerating microspores starting in anthers containing early microspores. (c) Anti-GFP staining of male meiosis stages in a *proSKR:GFP-gSKR* complementing line; asterisks: somatic/tapetal cells; arrowheads: older uninucleate microspores with low GFP signal. Scale bar: (a) 5 µm, (b,c) 20 µm.

### Early male gametogenesis defects in the *shukr* mutant

Analysis of meiosis and pollen development was performed to identify the stage at which defects arose in *skr*. Pollen development is divided into two stages: microsporogenesis comprising meiosis together with microspore development, and microgametogenesis or male gametophyte development comprising development of the haploid microspore into pollen^60^. No defects were observed in *skr* during meiosis based on analysis of chromosome spreads (Fig. 2a). Light microscopic examination of whole mount cleared anthers and plastic sections, showed that tetrad formation and release of microspores took place in *skr* (Figs. 1h,2b; Extended Data Fig. 3), however, microspores degenerated shortly after release from tetrads (Fig. 2b; Extended Data Fig. 4). Newly released wild type microspores were angular in shape (Fig. 2b)^61^ whereas *skr* microspores were more rounded and variable in cell wall thickness (Fig. 2b), suggesting loss of rigidity and possible defects in cell wall development. Hence sterility in *skr* appeared to be associated with defects in male gametogenesis between tetrad formation and early gametophyte development.

Ultrastructural examination of tetrad and early microgametogenesis stages (Fig. 3a) by transmission electron microscopy revealed that a significant proportion of *skr* tetrads harbored microspores in which the plasma membrane (PM) was retracted from the surrounding callose wall (Fig. 3b-d). PM retraction was associated with presence of material between the PM and callose layer likely reflecting unincorporated cell wall precursors or disrupted material arising from disturbance in cell wall development. Hence onset of the mutant phenotype was detected at the tetrad stage. To test for additional defects in *skr*, we examined expression of the LAT52:GFP reporter that specifically marks the single vegetative cell after pollen mitosis 1^62^. Among *skr* microspores in which expression of LAT52:GFP took place, we observed two signals in 30% of cases (Fig. 3e). Therefore loss of *SKR* led to at least two distinct defects in early gametogenesis: retraction of the PM with associated disruption of cell wall development, and aberrant expression of an early differentiation marker.

**Fig. 3:**
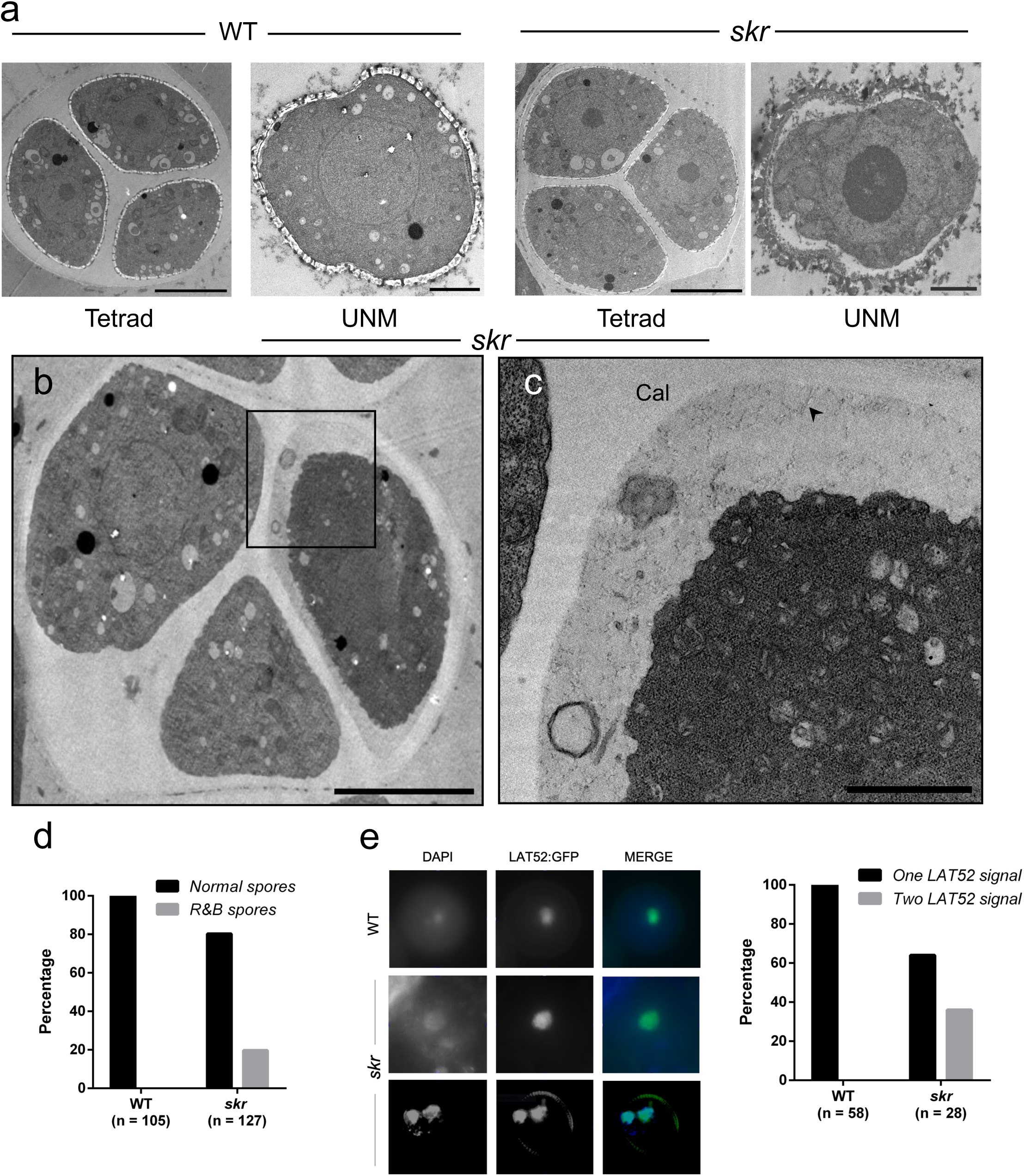
Ultrastructure and marker expression reveal early gametogenesis defects in *skr*. (a-c) Ultrastructure of tetrad and early uninucleate microspores showing onset of mutant phenotype at tetrad stage in *skr*. (a) WT: Normal tetrad and microspore; *skr*: normal tetrad, degenerating microspore with retracted PM. (b) *skr* tetrad in which one spore shows PM retraction from callose wall. (c) Magnified image of boxed region from (b) showing PM retraction from callose wall (Cal) and disrupted cell wall material (arrowhead). (d) Quantitation of normal spores and spores showing PM retraction and broken cell wall material (R&B) in tetrads (Fisher’s exact test p < 0.001). (e) Presence of two LAT52:GFP signals in a proportion of *skr* microspores (Fisher’s exact test p < 0.001). Scale bar (a): Tetrad images 5 µm; microspore images 2 µm; (b) 5 µm; (C) 1 µm.

Recessive gene action in the diploid sporophyte would give Mendelian 3:1 segregation of the mutant allele. Gene requirement in the haploid male gametophyte would result in reduced transmission of the mutant allele through the male and a deviation from this ratio. The genetic analysis suggested that *SKR* acts sporophytically as we did not observe a significant deviation from this ratio (Fig. 4a), whereas the mutant phenotype is with respect to the early gametophytic phase. We tested whether *SKR* also has a gametophytic component to its action by reciprocal crosses of *skr/+* to wild type and examining transmission frequency of the mutant *skr* allele through the gametophyte in comparison to the wild type allele. This test measures fitness of mutant relative to wild type gametes from the same flowers. If there is a functional requirement for *SKR* gene expression at the haploid stage in the gametophyte, then mutant pollen would show reduced fitness and the mutant allele would be transmitted to the next generation at lower frequency than the wild type allele. The results indicated no significant difference in transmission frequency of the mutant *skr* allele from that of the wild type (Fig. 4a), thus failing to provide evidence for *SKR* gene action within the gametophyte. Therefore the genetic results together with ultrastructure and marker expression indicated that *SKR* gene action is in the diploid sporophyte and is required for proper execution of early stages of gametogenesis. Further tests confirmed this finding (see below).

**Fig. 4:**
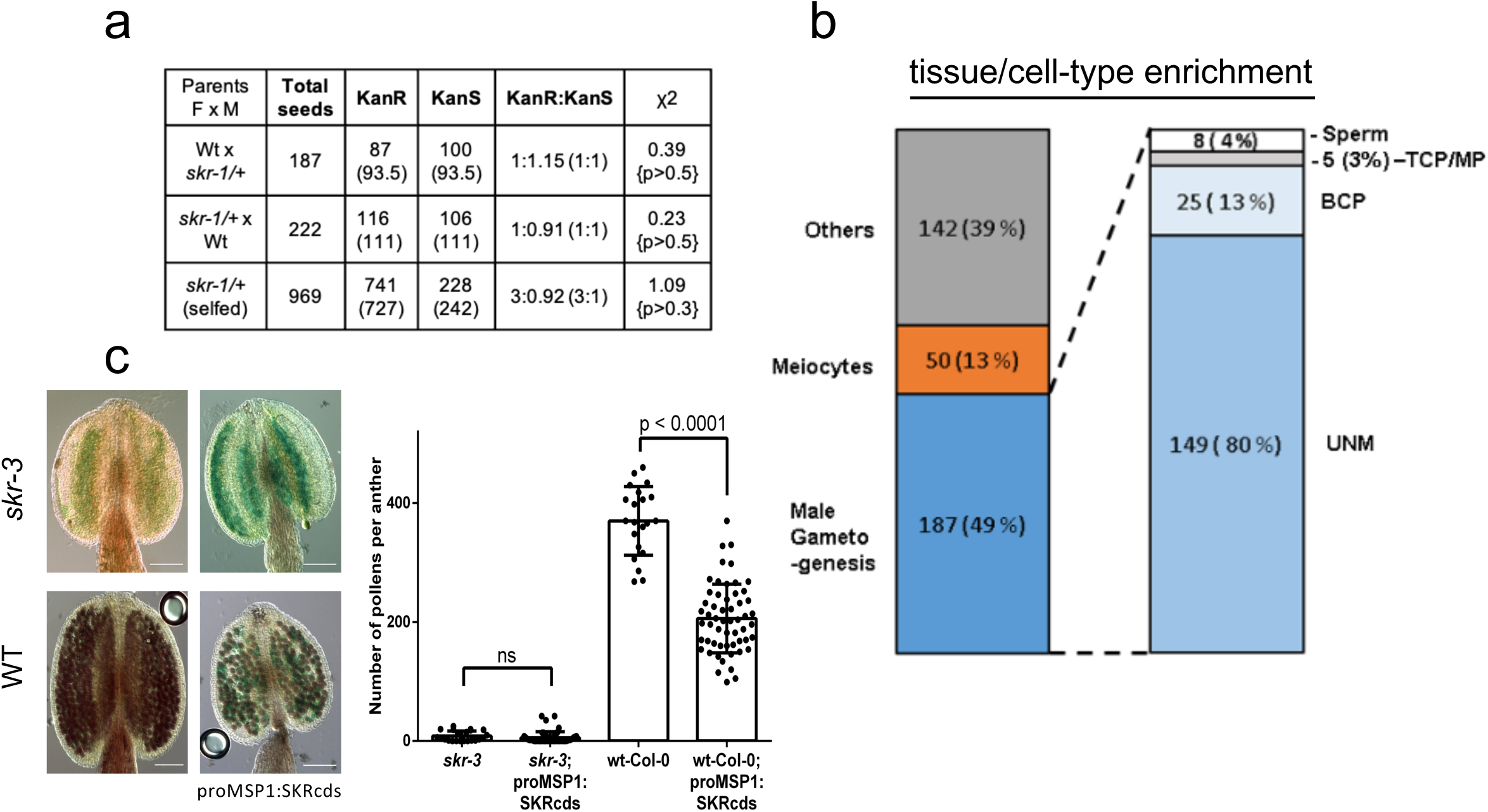
*SHUKR* is a gametogenesis inhibitor. (a) Reciprocal crosses comparing transmission of *skr* and wild type alleles; numbers in brackets are those expected for equal transmission and χ^2^ test p-value. (b) Tissue/cell enrichment categories for upregulated genes in *skr*. (c) Pollen viability in WT and *skr* plants transgenic for *proMSP1:SKRcds* or control. Data are pools of 4-10 anthers from 5 independent transgenic lines in each case. Scale bar (c) 100 µm.

### Meiotic control of male gametophyte gene expression by *SHUKR*

The sporophyte is known to play an important role in formation of the pollen wall and aperture^60,63^. Much of the material and machinery for building the outer wall (exine), is synthesized and released by the tapetum, a layer of nurse cells adjacent to the meiocytes^47,64^. Examination of *SKR* gene expression described above indicated that *SKR* is expressed specifically in male meiocytes. Therefore unlike the sporophytic genes which function in the tapetum, *SKR* appears to act in meiosis. We determined the localization of SKR protein in the course of meiosis using a complemented line carrying a *proSKR:GFP-gSKR* gene fusion. SKR protein was detected specifically in male meiocytes from prophase to tetrad stage and extending to early microspores. Subcellular localization of GFP-SKR varied during meiosis, from nuclear enrichment at prophase, changing toward cytoplasmic in late prophase, and back to nuclear at the tetrad stage. Signal intensity was highest in tetrads (Fig. 2c). Localization in early microspores shifted from nuclear towards more of cytoplasmic, before declining to an undetected level in later stage uninucleate microspores (UNM). Expression was not detected in tapetal cells (Fig. 2c). Therefore SKR protein is mainly present during meiosis and the onset of the *skr* mutant phenotype corresponds to the tetrad stage at which SKR signal is highest. High SKR expression in tetrads followed by steep decline in free microspores suggests a possible role in repressing gametogenesis or else in ending the sporophytic program before initiation of gametogenesis.

To investigate the above possibilities we examined gene expression during meiosis by RNAseq profiling of meiotic anthers (extending upto and not beyond tetrad stage) for *skr* and wild type. 619 genes were differentially expressed in *skr* of which 379 genes were upregulated (Supplementary Table 1). We analyzed upregulated genes based on cell and tissue expression in publicly available datasets using the CoNekT platform^65^ and found a dramatic enrichment for male gametogenesis-enriched genes^66,67^ (Fig. 4b, Supplementary Table 2). 187/379 (49%) of upregulated genes were male gametogenesis-enriched. Genes showing maximal enrichment in UNMs represented 80%, followed by bicellular and tricellular pollen stages (16%). Upregulated genes included *DML3* which has been suggested to be an early gametogenesis marker^68^. Increased expression of a large set of gametogenesis-enriched genes in meiotic anthers of *skr* suggests that SKR negatively regulates expression of gametogenesis genes and represses gametogenesis.

### SHUKR acts sporophytically and inhibits gametogenesis

To test for a role in repression of gametogenesis as well as the possibility that SKR protein acts postmeiotically to promote pollen development, we adopted a misexpression strategy by directing expression of SKR in microspores using the well characterized microspore specific MSP1 promoter^69^. *proMSP1:SKR* in a *skr* mutant failed to rescue the mutant phenotype, indicating that SKR expression in the gametophyte is not sufficient to restore function. Furthermore, *proMSP1:SKR* in wild type in fact caused a reduction in pollen viability (Fig. 4c). These results show that expression of SKR during gametogenesis interferes with gametophyte development and argue against the possibility that SKR protein present during postmeiotic stages contributes to pollen development. Therefore SKR acts sporophytically and is an inhibitor of male gametogenesis.

### *SHUKR* represses the ubiquitin proteasome system

Gene ontology (GO)^70^ together with further analysis of individual gene families revealed a striking signature of regulated protein turnover among the 379 genes upregulated in *skr* RNAseq. 23% of upregulated genes were associated with ubiquitin-mediated degradation of proteins^71^ (Fig. 5a, Supplementary Table 3; p < 2.0e-33). At the level of molecular function, there was enrichment for ubiquitin protein transferase activity covering the Skp1-Cullin-Fbox (SCF) complex of E3 ubiquitin ligase as well as other components of the ubiquitin protease system (UPS) including *RING/U-box* and *ARIADNE* genes which encode E3 ubiquitin ligases^72,73^. F-box genes encode the substrate specificity component of SCF and comprised the largest family among upregulated genes in *skr*. F-box genes in Arabidopsis make up a family of approximately 740 genes^74,75^ (Supplementary Table 3), several of which show expression during male gametogenesis^76^. 69 F-box genes representing 9% of all F-box genes in Arabidopsis were upregulated in *skr* (R.F. = 6.6, p < 1.74e-36; Fig. 5a; Supplementary Table 3). Hence a significant fraction of F-box genes are implicated in pollen development. 75% of upregulated F-box genes were maximally enriched at the UNM stage and 9% at later stages comprising bicellular and tricellular pollen. Among downregulated genes, there was enrichment for stress and hypoxia related genes (Supplementary Table 3). Upregulation of a large number of UPS genes suggested involvement of SKR in general control of protein homeostasis and further experiments were performed to test this hypothesis (see below). We also examined stage-wise expression of a subset of upregulated UPS genes at the cellular level, by generating lines carrying a GFP gene fusion for three different F-box genes (*At3g28330*, *At5g02980* and *At5g62510*) and segregating for *skr*. In each case, we observed F-box:GFP expression in wild type only during postmeiotic development of microspores into pollen, whereas *skr* showed precocious F-box:GFP expression in meiosis (Fig. 5b, Extended Data Fig. 5). Hence SKR can be seen to act in meiosis to bring about repression of these F-box genes. Taken together, these observations led us to hypothesize that SKR represses onset of male gametogenesis during meiosis through timing of expression of a large set of male gametophyte-preferred F-box and other UPS genes.

**Fig. 5:**
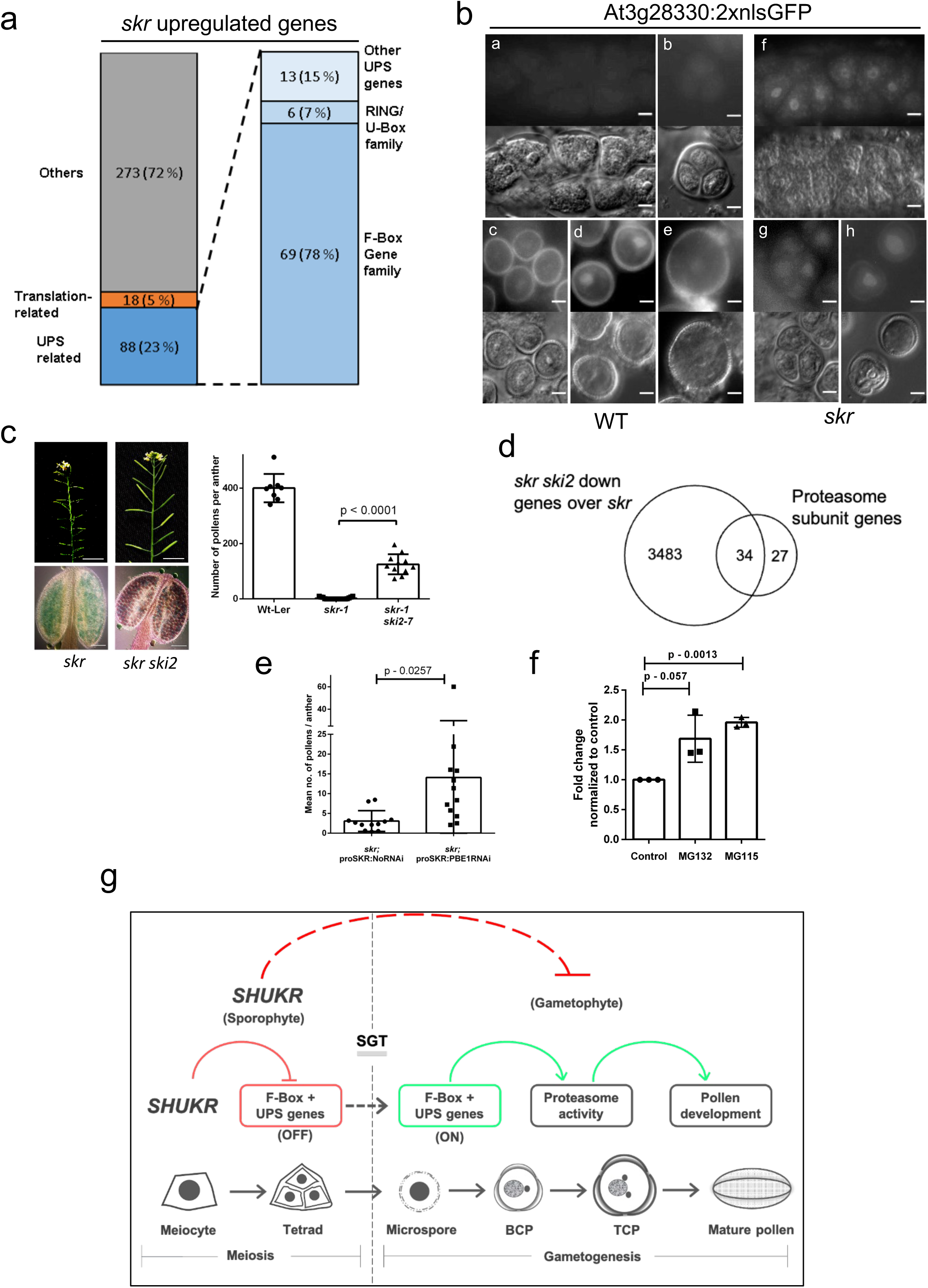
*SHUKR* regulates the proteasome in pollen development. (a) Gene family categories for upregulated genes in *skr*. (b) Stagewise expression of an *At3g28330:2xnlsGFP* transgene during meiosis and in microspores for *skr* and wildtype. Frames: a,f meiocytes; b,g tetrad; c-e,h microspores; upper half: GFP, lower half: DIC. (c) Suppression of *skr* male sterility by *ski2*. Bar graph shows viable pollen counts in single anthers from 2-4 plants. (d) Downregulation of proteasome genes in *skr ski2*. (e) Mean viable pollen counts/anther in independent *skr; proSKR:PBE1RNAi* and control *T1* transgenic plants. (f) Pollen viability increase in *skr* following proteasome inhibitor treatment in 3 replicate experiments. (g) Model for sporophytic control of pollen development by *SHUKR*. During meiosis *SHUKR* establishes a programme that determines timing of expression of F-box and other UPS genes following meiosis. Scale bar: (b) 5 µm; (c) plant 1 cm, anthers 100 µm. Abbreviations: UNM, uninucleate microspore; BCP, bicellular pollen; TCP, tricellular pollen; MP, mature pollen; SGT, sporophyte-gametophyte transition.

### Control of protein homeostasis by *SHUKR*

We directly investigated the functional basis for the *skr* phenotype by performing a suppressor screen on *skr* using EMS mutagenesis of seeds collected from a large number of *skr* plants followed by screening of M2 plants for improved seed set. We isolated a suppressor of *skr* which showed elongated siliques and increased pollen viability, whereas *skr* shows low pollen viability. The suppressor line was backcrossed to the parent *skr* strain and F1 plants were sterile. F2 seed were collected from multiple F1 plants and grown to give F2 plants which showed recessive Mendelian inheritance for fertility (Extended Data Fig. 6a). Whole genome sequencing of bulk segregant fertile and sterile plants from the backcrossed F2 population led to identification of an SNP at the *SKI2* locus which created a stop codon after 517 amino acids and showed 100% linkage to the suppressor phenotype (Fig. 5c, Extended Data Fig. 6b). *SKI2* encodes the RNA helicase subunit of the SKI complex and prevents secondary siRNA production in Arabidopsis^77^. *ski2* has also been isolated as a suppressor of overexpression phenotypes in a different context^78^. We tested an independent allele of *ski2* by crossing to *skr* and observed suppression of the *skr* mutant phenotype, confirming *ski2* as a suppressor of *skr* (Extended Data Fig. 6c). We then performed RNAseq on meiotic anthers of the *skr ski2* backcrossed suppressor line. Comparison of differentially expressed genes with the *skr* parent revealed a prominent signature of the proteasome and translation among downregulated genes: 56% of genes encoding subunits of the proteasome and 29% of translation related genes were down-regulated in *skr ski2* (Fig. 5d, Supplementary Table 4). Notably, F-box genes as a class were not downregulated in *skr ski2* (R.F.= 0.3; p < 7.7e-19). The striking reduction in transcript levels of the majority of proteasomal genes in *skr ski2* strongly supports the hypothesis that the *skr* phenotype arises from uncoordinated degradation by the proteasome, of regulator(s) of gametogenesis due to inappropriate expression of gametogenesis specific F-box and possibly other ubiquitination genes. Reduction of proteasome activity by downregulation of a major subset of proteasome genes rescues the mutant phenotype.

We confirmed involvement of the proteasome in the *skr* phenotype using two different approaches. First, we performed knockdown of PBE1, a core component of the proteasome^79^ in the *skr* mutant background using *pro*SKR to direct meiosis-specific expression of PBE1 hairpin RNA and scored for pollen viability in RNAi plants. Knockdown of PBE1 by RNAi has been used in a different context^80^ and we used the same region for hairpin RNA construction in our experiments. Second, we treated inflorescences of *skr* mutant plants with two physiological inhibitors of the proteasome (MG132 and MG115). In both regimes, we observed an increase in formation of viable pollen that was significant for PBE1:RNAi and MG115 (Fig. 5e,f; p < 0.05). Taken together, these results clearly establish deregulation of proteasome activity as a major basis for the *skr* mutant phenotype. The large number of F-box genes showing deregulation during meiosis in *skr*, most of which normally are preferentially expressed only after meiosis during gametogenesis, highlights the scope of this control. Hence, SKR plays a crucial role in meiosis to repress a proteasomal program that promotes gametogenesis and which comes into play in free microspores following meiosis. SKR therefore sporophytically regulates postmeiotic development of the gametophyte through timing of regulated protein turnover by the proteasome (Fig. 5g). Notably, this programme is distinct from the canonical meiotic processes of chromosome reduction and spore formation, which remain unaffected in *skr*.

### *SHUKR* is eudicot specific and rapidly evolving

A cDNA of the *SKR* coding region was amplified by RT-PCR, sequenced, and found to encode a predicted protein of 191 amino acids in agreement with the annotated CDS. Homologs were found only in plants and bore no similarity to any known protein family. SHUKR is therefore the founding member of a new protein family (hereinafter SHUKR). The only alignable region conserved across all SHUKR homologs is always at the C-terminus and this is usually the only globular domain in the protein from Shannon entropy-based compositional analysis. The domain has a central compositionally biased, poorly structured variable region enriched in negatively charged residues. Thus, the SHUKR domain varies in length from 67 to 119 residues. It has a hallmark CxC motif and a dyad of aromatic amino acids (usually Y or F) close to the end of the domain (Fig. 6a, Supplementary Figure 1). The SHUKR family is present in both great divisions of eudicots: the Superrosids and Superasterids. It is additionally present in Proteales (Nelumbo) and Ranunculales (Macleaya, Papaver), which are the two basal-most branches of eudicots. It is absent in monocots and basal angiosperms. Hence the origin of SHUKR is at the base of eudicots.

**Fig. 6:**
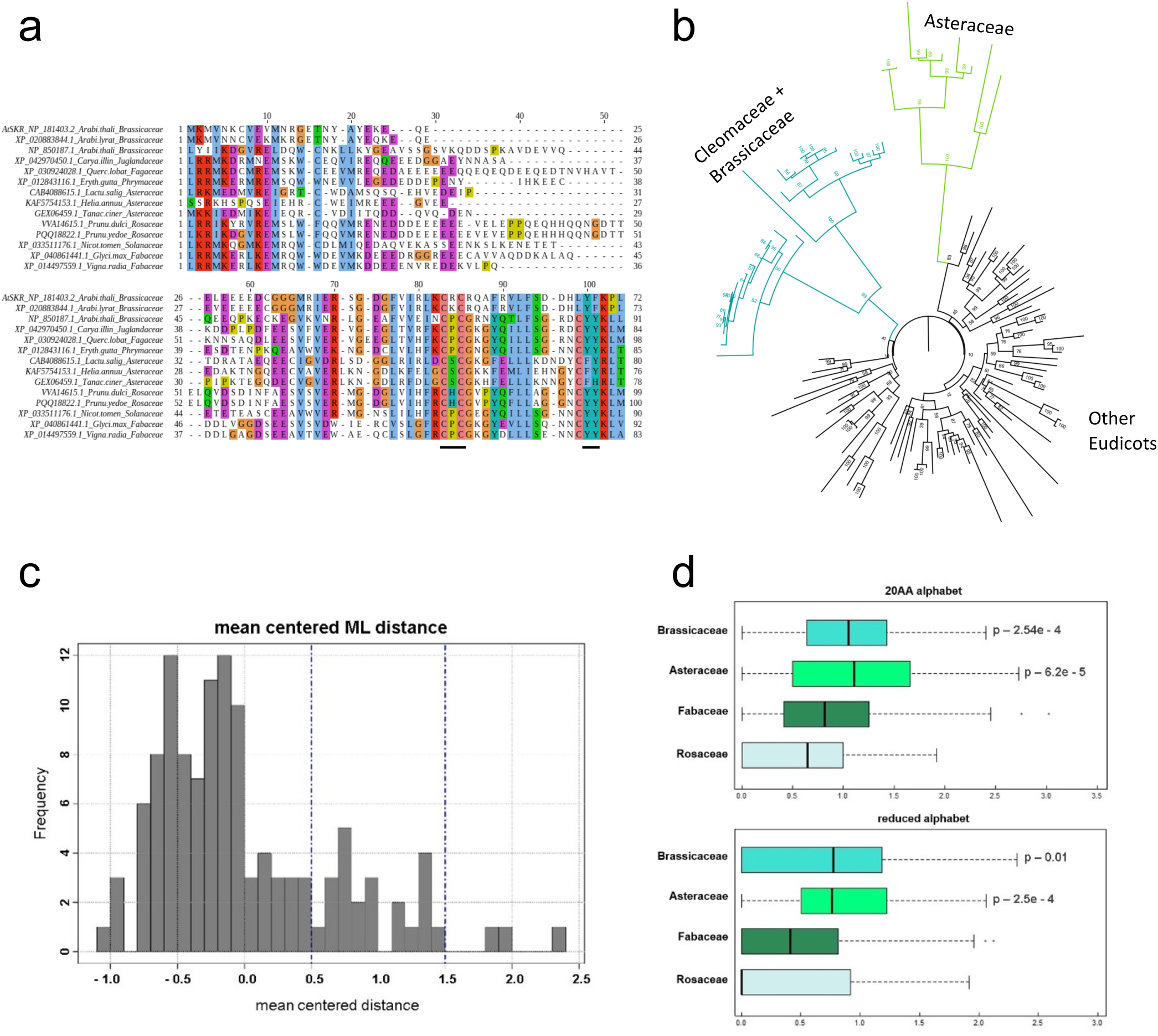
Sequence and evolutionary analysis of the SHUKR domain. (a) Multiple sequence alignment of the SHUKR domain. Lines mark CxC and aromatic dyad. (b) Phylogenetic tree of the SHUKR domain from 119 species. Dark green, Brassicaceae+Cleomaceae; Light green, Asteraceae. (c) Distribution of mean-centered Maximum Likelihood (ML) distance differences. (d) Columnwise Shannon entropy distributions for four plant families. P-value calculated using Kruskal-Wallis test for Brassicaceae and Asteraceae relative to the other two families.

To understand evolution of the SHUKR family we used the SHUKR domain of 119 representatives from 85 species for tree construction. Branch length analysis of the tree revealed two lineages displaying elevated evolutionary rates: Brassicaceae+Cleomaceae and Asteraceae (Fig. 6b; Extended Data Fig. 7). This was further tested using the maximum-likelihood distance matrix computed from the input alignment for the tree. The mean pairwise distance was calculated between each operational taxonomic unit (OTU) and rest of the OTUs, and centered using overall mean distance for the matrix. These mean-centered distances showed a bimodal distribution with the main mode between −1.25 to 0.5, a second mode from 0.5 to 1.5, and a few outliers beyond that (Fig. 6c). The second mode corresponds to sequences with higher values of mean-centered distance, indicating their more extensive divergence. Brassicaceae+Cleomaceae (p = 1.1472e-04) and Asteraceae (p = 1.6e-16) were significantly enriched among sequences with mean-centered distance greater than 0.5 indicating they represent atypically fast-evolving groups. We note that the high divergence of the SHUKR family outside the SHUKR domain posed challenges in alignment across the entire coding sequence. Therefore the analysis likely represents a conservative assessment of evolutionary rates within the *SHUKR* gene family.

### *SHUKR* is under positive selection

We assessed rapid evolution in particular plant clades by computing column-wise Shannon entropy from an alignment of the SHUKR domain for four plant families with good representation in the Genbank nr database: Brassicaceae, Asteraceae, Fabaceae and Rosaceae. The column-wise entropy was computed using both the 20 amino acid alphabet and a reduced 7 amino acid alphabet that groups together residues as per their shared sidechain chemistry (see Methods). Elevation of entropy in the reduced alphabet indicates changes in sidechain properties along a column and is a proxy for diversifying selection^81^. Entropy plots in both alphabets indicated significantly elevated entropy in Brassicaceae and Asteraceae relative to other families, suggesting action of diversifying selection in these lineages (Fig. 6d). At least 30% of species examined had more than one paralog per genome most of which appeared to represent recent lineage-specific duplications as they tended to group together in the tree. However, Asteraceae appears to have experienced a duplication early in their history. Likewise the duplication in Brassicaceae and Cleomaceae predated their divergence. We further assessed diversifying selection at *SHUKR* within Brassicaceae based on an alignment spanning the length of the gene, using branch-site models^82^ that consider proteins as a mosaic of conserved and divergent domains,. The branch-site test results provided further support for existence of positive selection (p = 0.0031) (Extended Data Table 1; Extended Data Fig. 8; Supplementary Figure 2), demonstrating that gene duplication has been followed by diversifying selection at *SHUKR*.

### Adaptive value for variation in protein turnover during male gametogenesis

Plant F-box genes have undergone independent lineage-specific expansions as well as reductions during land plant evolution^83,84^. The Brassicaceae family has an expanded set of F-box genes, a majority of which are of short taxonomic scale indicative of more recent emergence^84^. These recent members were significantly enriched among upregulated F-box genes in *skr* (p < 1.5e-05) (Fig. 7a,c; Supplementary Table 5). Orthology analysis of the total set of F-box genes from 10 eudicot species^84^ indicated that 58% (37/64) of F-box genes upregulated in *skr* belonged to orthogroups that included *Arabidopsis lyrata* but not 8 other eudicot species examined spanning 7 families including *Carica papaya* (family *Caricaceae*, order *Brassicales*), suggesting that these were likely specific to the Brassicaceae lineage (Fig. 7b; Supplementary Table 5). 8% (5/64) of genes belonged to orthogroups having only *A. thaliana* and not *A. lyrata* suggesting a more recent origin, and 23% (15/64) had at least one other species but not *A. lyrata* indicative of recent loss (Fig. 7b). We examined the possibility of diversifying selection at the set of *skr* upregulated genes by testing whether the ratio K_a_/K_s_ of non-synonymous to synonymous substitutions per site was higher relative to the overall set of F-box genes. Plots of K_a_/K_s_ gave significantly higher values for the set of F-box genes upregulated in *skr* relative to the overall set of F-box genes (p = 0.0075; Fig. 7d). Hence the set of *skr* upregulated F-box is itself undergoing diversifying selection. Interestingly, male gametes have been proposed as a source for emergence of new genes in both animals and plants^85–87^, and pollen-preferred F-box genes also appear to belong to this set. Variation in protein turnover during male gametophyte development can therefore be seen to operate at two levels in evolution. One is through rapid evolution and positive selection in a regulator of UPS genes involved in male gametophyte development. The second is in changes and diversifying selection in components of the UPS system itself, the set of F-box genes that direct substrates for proteasomal degradation in gametogenesis. The large number of F-box genes involved and their distribution across the F-box phylogenetic tree (Fig. 7c) suggests the likelihood of levels of multiple proteins being subject to regulation by SKR. Variation in this control provides a plausible route for the sporophyte to influence development and evolution of the gametophyte. These findings lead us to hypothesize that processes that provide for variation in levels of large sets of proteins rather than just a single target during male gametophyte development have adaptive value and have contributed significantly to evolution in Brassicaceae.

**Fig. 7:**
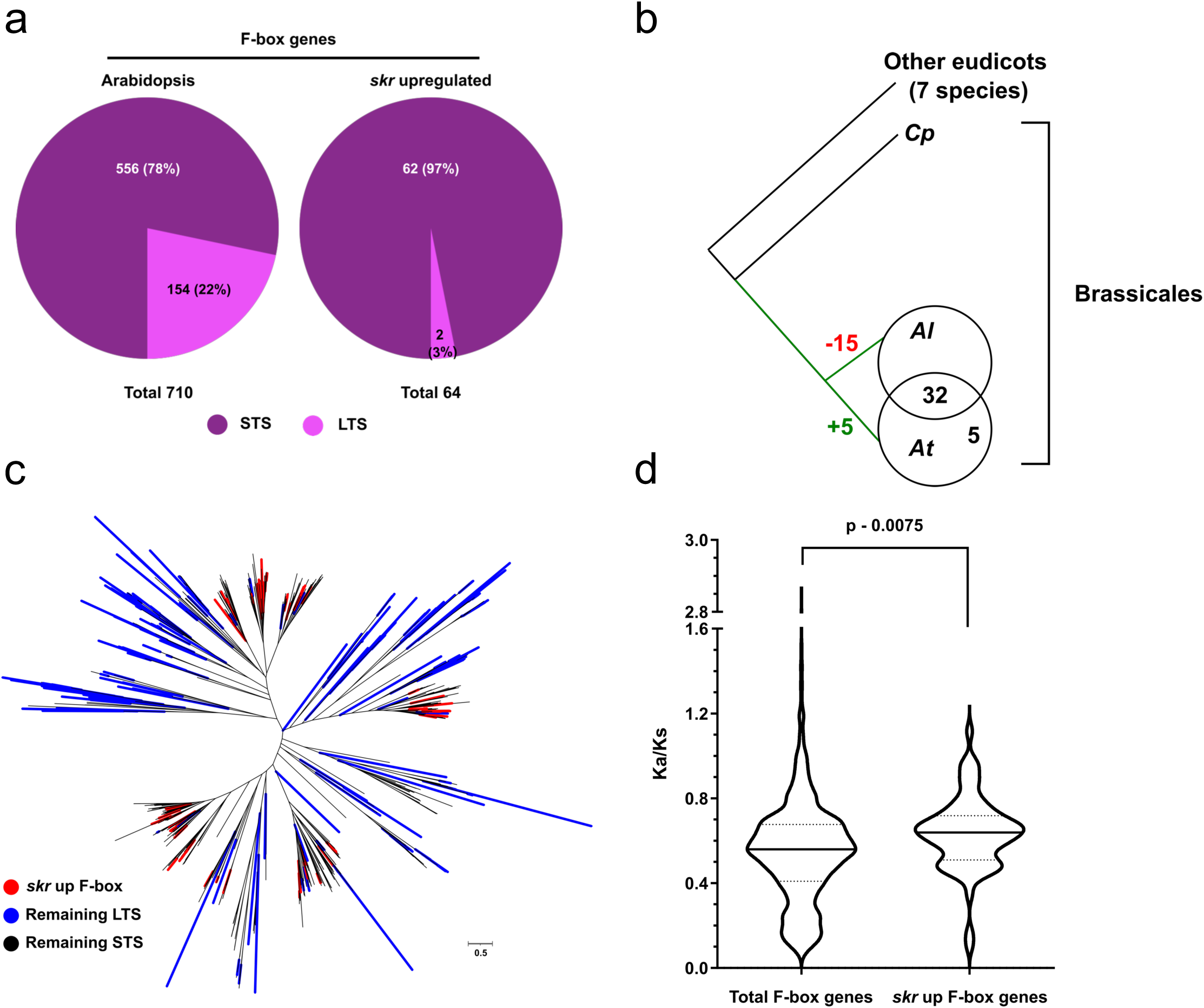
Diversifying selection in SHUKR regulated F-box genes. (a) Enrichment for short taxonomic scale (STS) F-box genes (R.F. = 1.2) and underenrichment for long taxonomic scale (LTS) genes (R.F. = 0.1) among *skr* upregulated set (p < 1.49e-05). (b) 37 genes were unique to the Brassicaceae lineage of which 5 were found only in *Arabidopsis thaliana*. 15 genes were present in *Arabidopsis thaliana* and at least one other species but absent in *Arabidopsis lyrata*. (c) Phylogenetic tree of Arabidopsis F-box genes showing *skr* upregulated, LTS, and STS sets. (d) Violin plot of K_a_/K_s_ for *skr* upregulated F-box and total F-box genes. Other eudicots: *Cucumis sativus*, *Glycine max*, *Manihot esculenta*, *Mimulus guttatus*, *Populus trichocarpa*, *Ricinus communis*, *Vitis vinifera*. Abbreviations: Al *Arabidopsis lyrata*, At *Arabidopsis thaliana*, Cp *Carica papaya*.

## Discussion

Extensive functional and gene expression studies of gametophyte development^15,18,88^ have supported the view that the male gametophyte retains control of its development even after evolutionary reduction to just a few cells enclosed within the sporophyte. Experiments in angiosperm species have also shown that isolated microspores can form pollen in culture^52,53,89^, supporting this view. Here we have demonstrated in Arabidopsis that the sporophyte in fact plays an important role in control of gametogenesis. This direction is by a repressive mechanism involving large scale control of regulated protein turnover in male gametophyte development by a novel gene, *SHUKR*. *SHUKR* acts during meiosis and is an inhibitor of gametogenesis (Fig. 5g). Our findings in Arabidopsis suggest that significant aspects of the male gametogenesis programme are established in the diploid sporophyte during meiosis but held in a repressed condition until microspores are released. We suggest the term sporophyte directed gametogenesis (SDG) for this phenomenon.

Recent emergence of *SHUKR* at the base of eudicots suggests that SDG arose late in evolution following reduction of the gametophyte. Gametophyte reduction and its close proximity to the sporophyte may have provided conditions necessary for the emergence of SDG. Rapid evolution and diversifying selection at *SHUKR* and the set of F-box genes under its control points to an adaptive value for variation in protein turnover in male gametophyte evolution. This variation provides a way for the sporophyte to generate diversity in the male gametophytes which face intense selective pressure in the course of transmission and intraspecific competition^90^. The location of this control in the sporophyte is also of interest because it raises the possibility that gametophyte development can be modulated by the sporophyte as a function of its own status and environment.

## Extended Data Figure Legends

**Extended Data Fig. 1:**
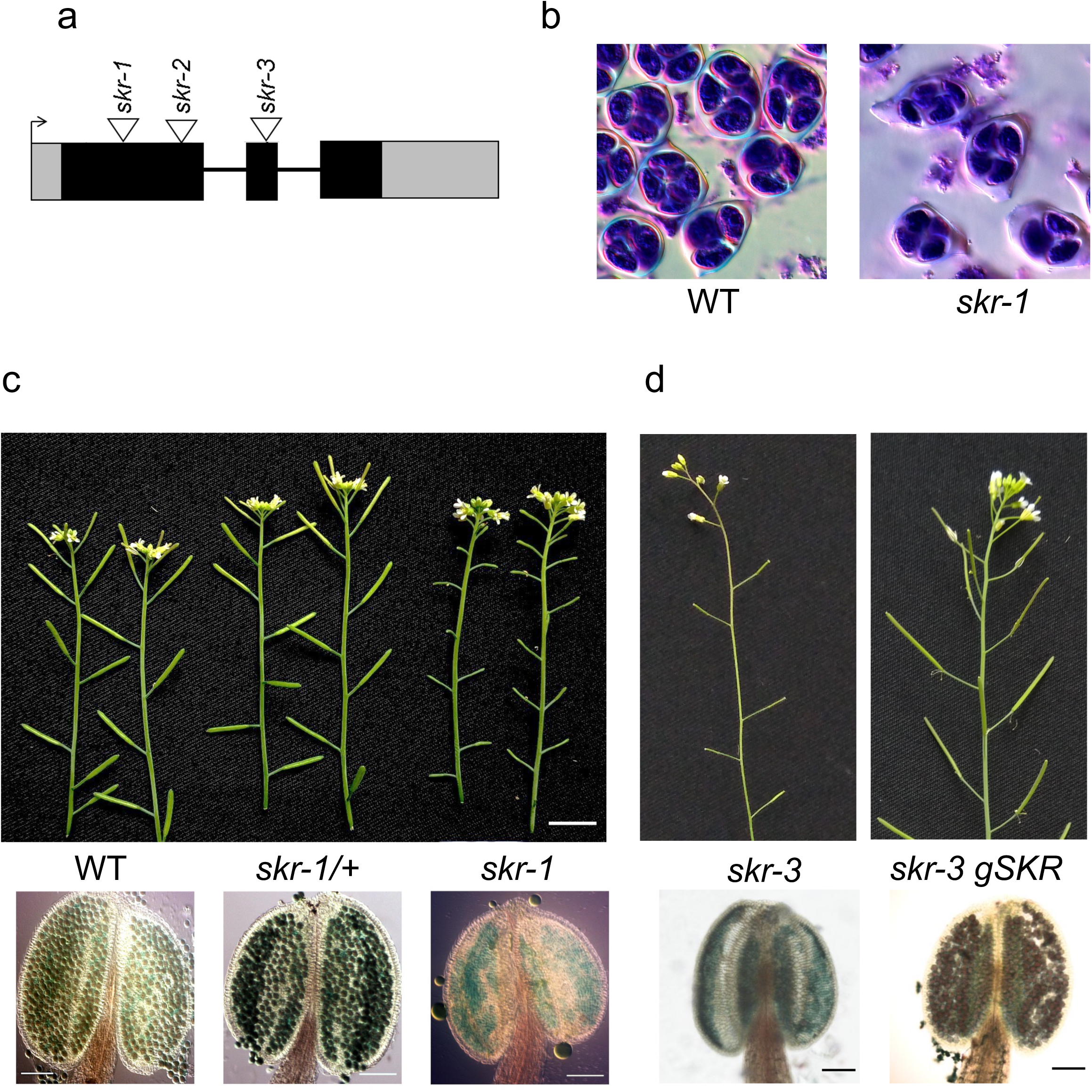
(a) Gene diagram of *SKR* showing location of insertions: *skr-1* (GT_5_101780), *skr-2* (CSHL_GT5061), *skr-3* (SALKSeq_034497.1). (b) Toluidine blue stained tetrads. (c) Inflorescence stem and Alexander staining showing recessive male sterility of *skr.* (d) Complementation of *skr-3* by a *gSKR* transgene. Scale bar: 1 cm for inflorescence, 100 µm for anthers.

**Extended Data Fig. 2:**
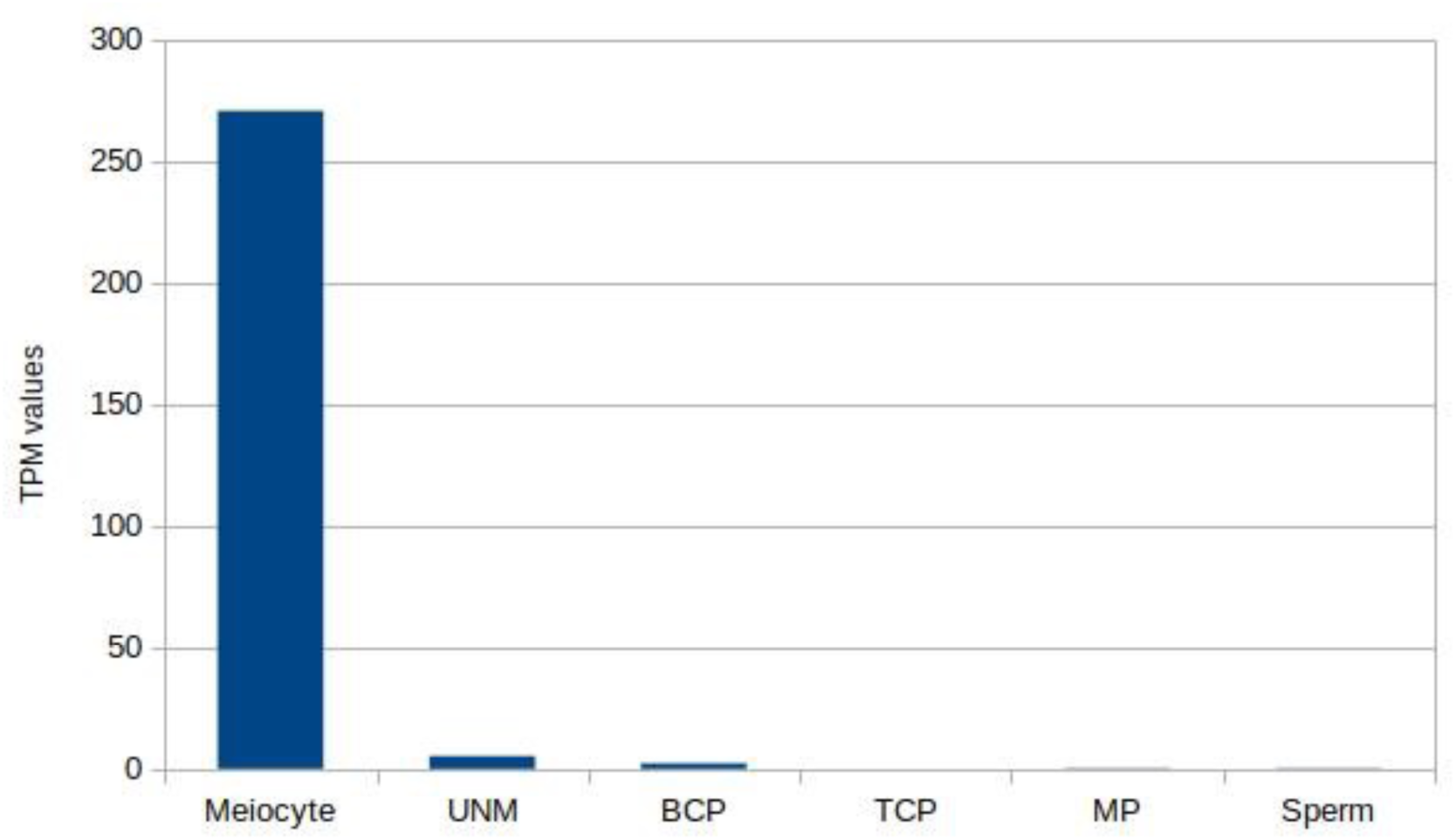
Profiling of SKR expression in male germline development. TPM values for SKR were collected from the CoNekt database, except for meiocytes. Meiocyte TPM value were taken from (Walker 2018)^21^. UNM – Uninucleate microspore; BCP – Bicellular pollen; TCP – Tricellular pollen; MP – Mature pollen.

**Extended Data Fig. 3:**
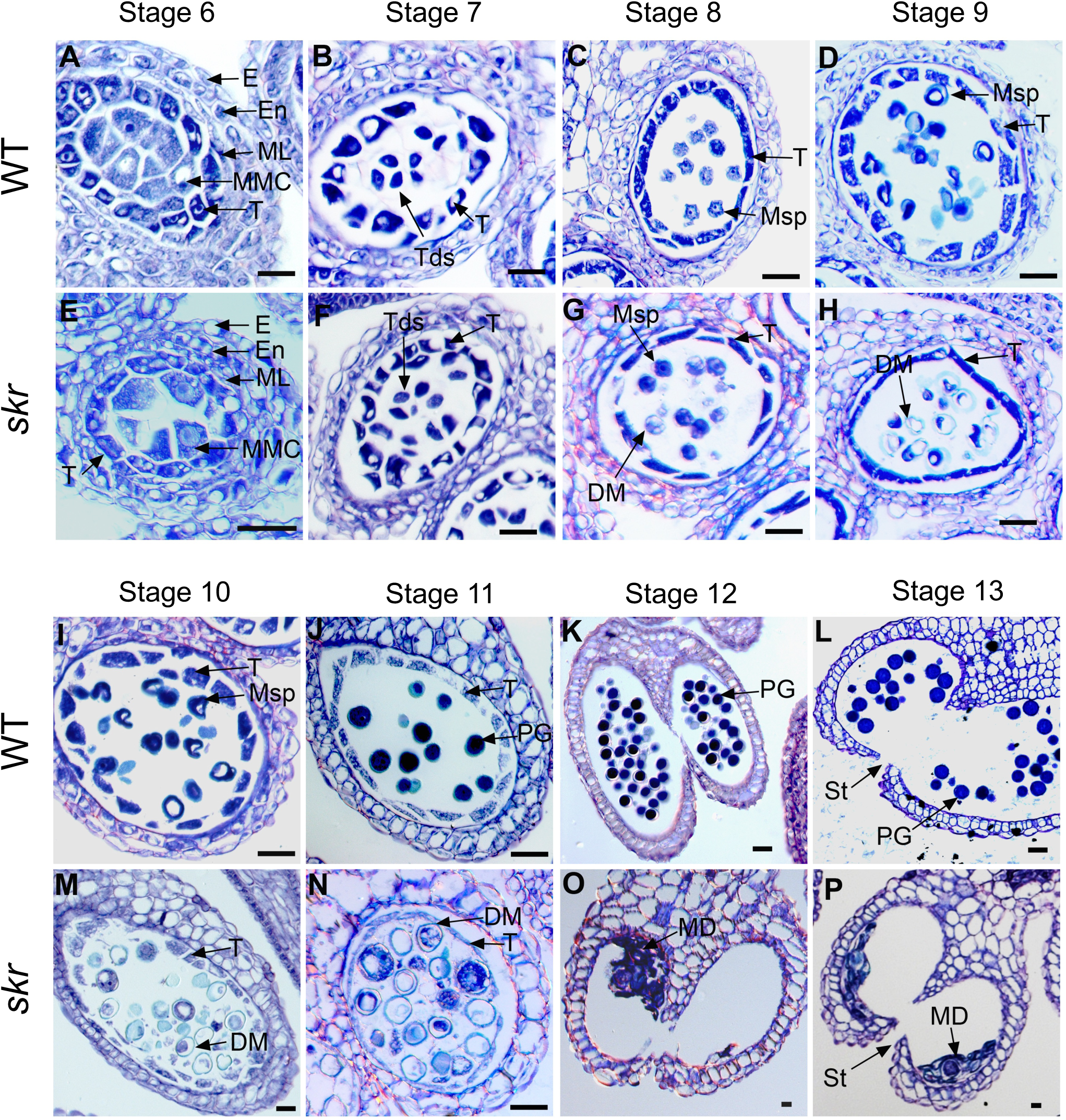
Sections of anther development stages. Corresponding stages are shown for wild type (A-D, I-L) and *skr* (E-H, M-P). Stages are numbered according to (Sanders 1999)^9^. Abbreviations: E epidermis, En endothecium, ML middle layer, T tapetum, Td tetrad, Msp microspore, DM degenerating microspore, PG pollen grain, St stomium, MD microspore debris. Scale bar 20 µm.

**Extended Data Fig. 4:**
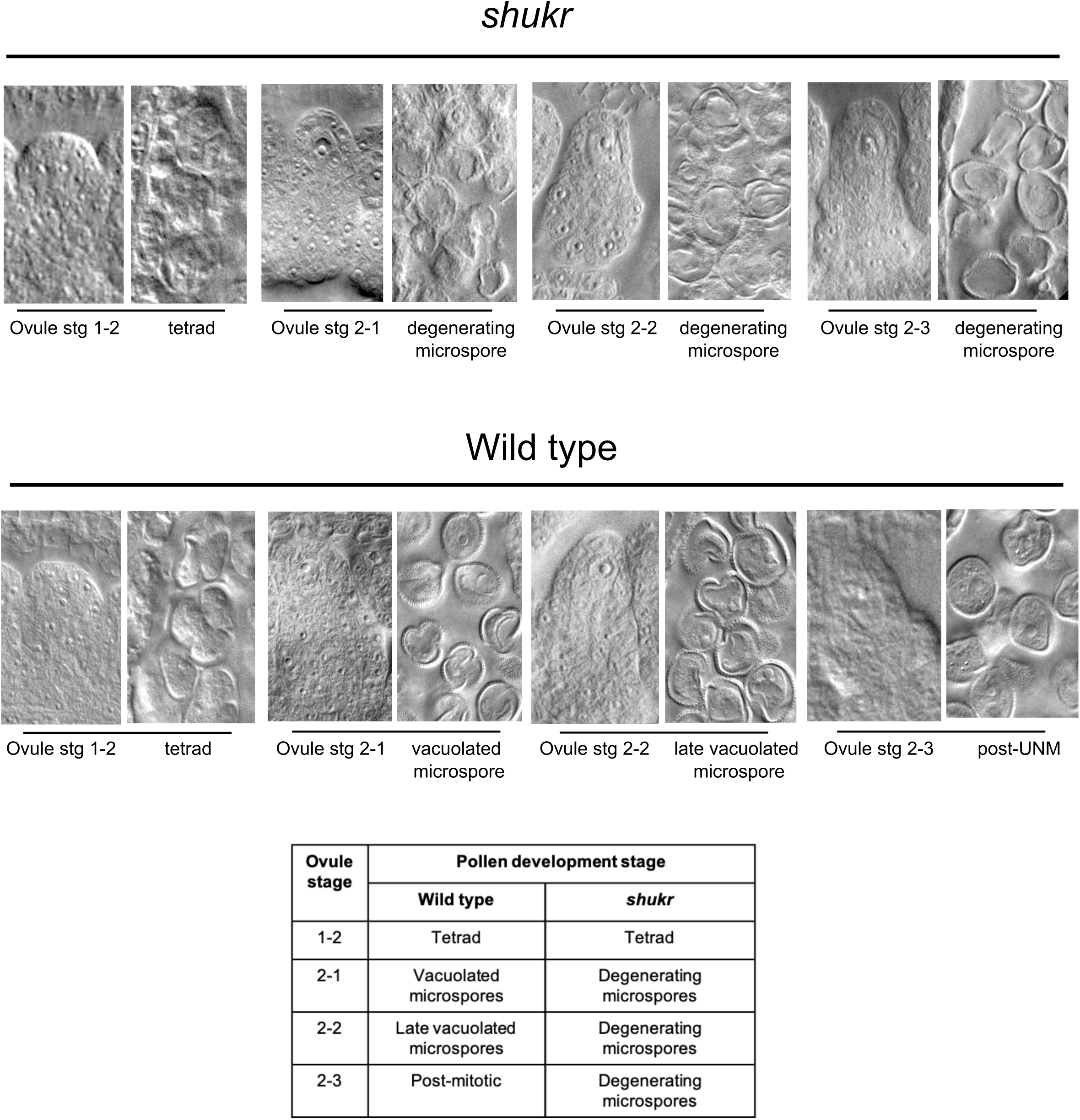
Staging of microspore degeneration in *skr*. Microspore stage and phenotypes are shown together with stage of ovules from the same bud (Schneitz 1995)^10^ as a reference for comparison to wild type

**Extended Data Fig. 5:**
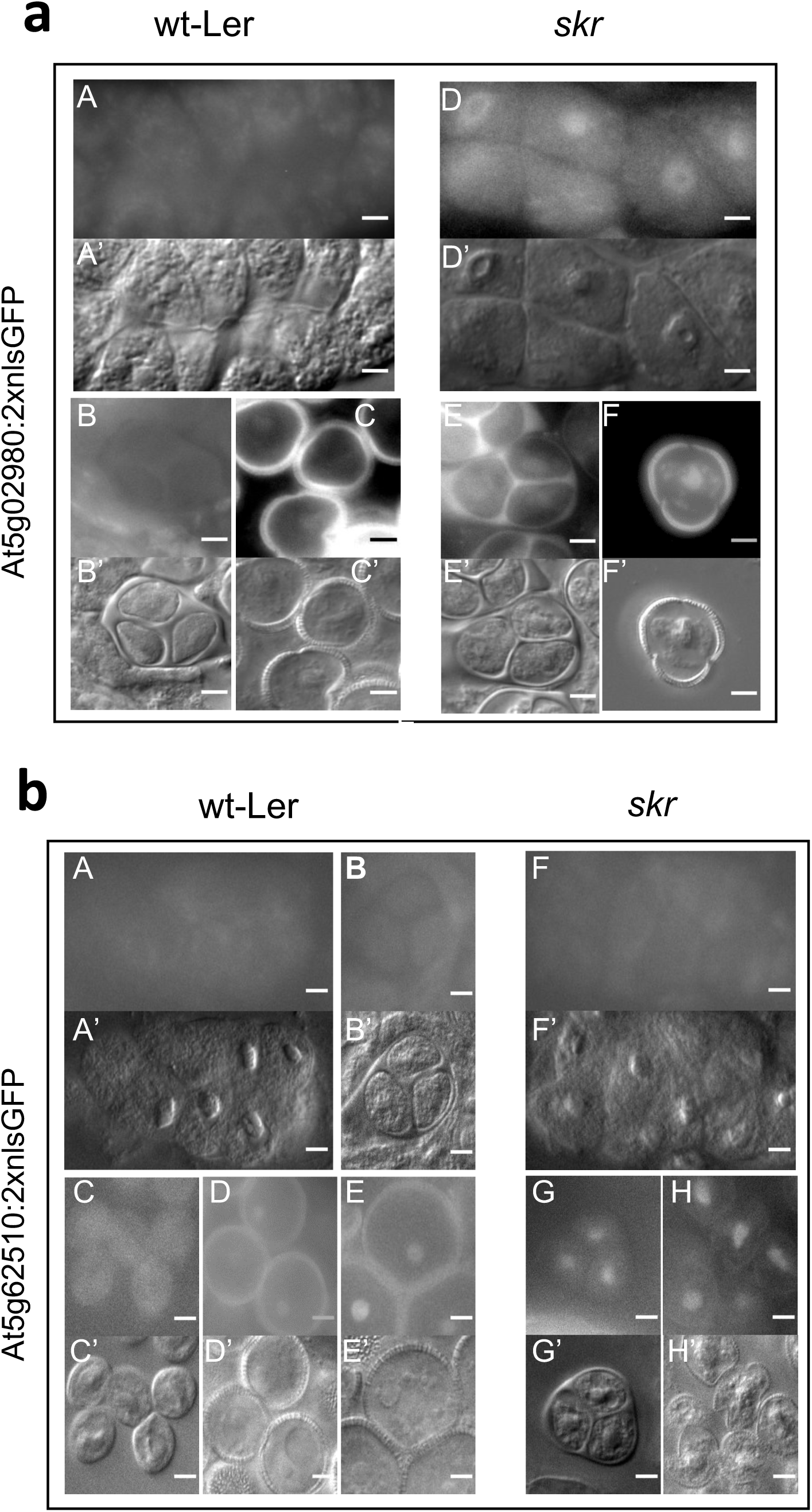
Precocious expression of F-box gene:nlsGFP fusions in meiosis in *skr*. (a) At5g02980. A-C, A’-C’: wild type; D-F, D’-F’: skr; A,A’,D,D’ meiocytes; B,B’,E,E’: tetrad; C,C’,F,F’: uninucleate microspores. (b) At5g62510. A-E, A’-E’: wild type; F-H, F’-H’: skr; A,A’,F,F’ meiocytes; B,B’,G,G’: tetrad; C-E, C’-E’,H,H’: uninucleate microspores. A-H: GFP fluorescence; A’-H’: corresponding DIC. GFP expression in *skr* is first detected in meiocytes (At5g02980) or tetrad stage At5g62510) whereas in wild type earliest GFP signal was in released microspores. Scale bar: 5 μm

**Extended Data Fig. 6:**
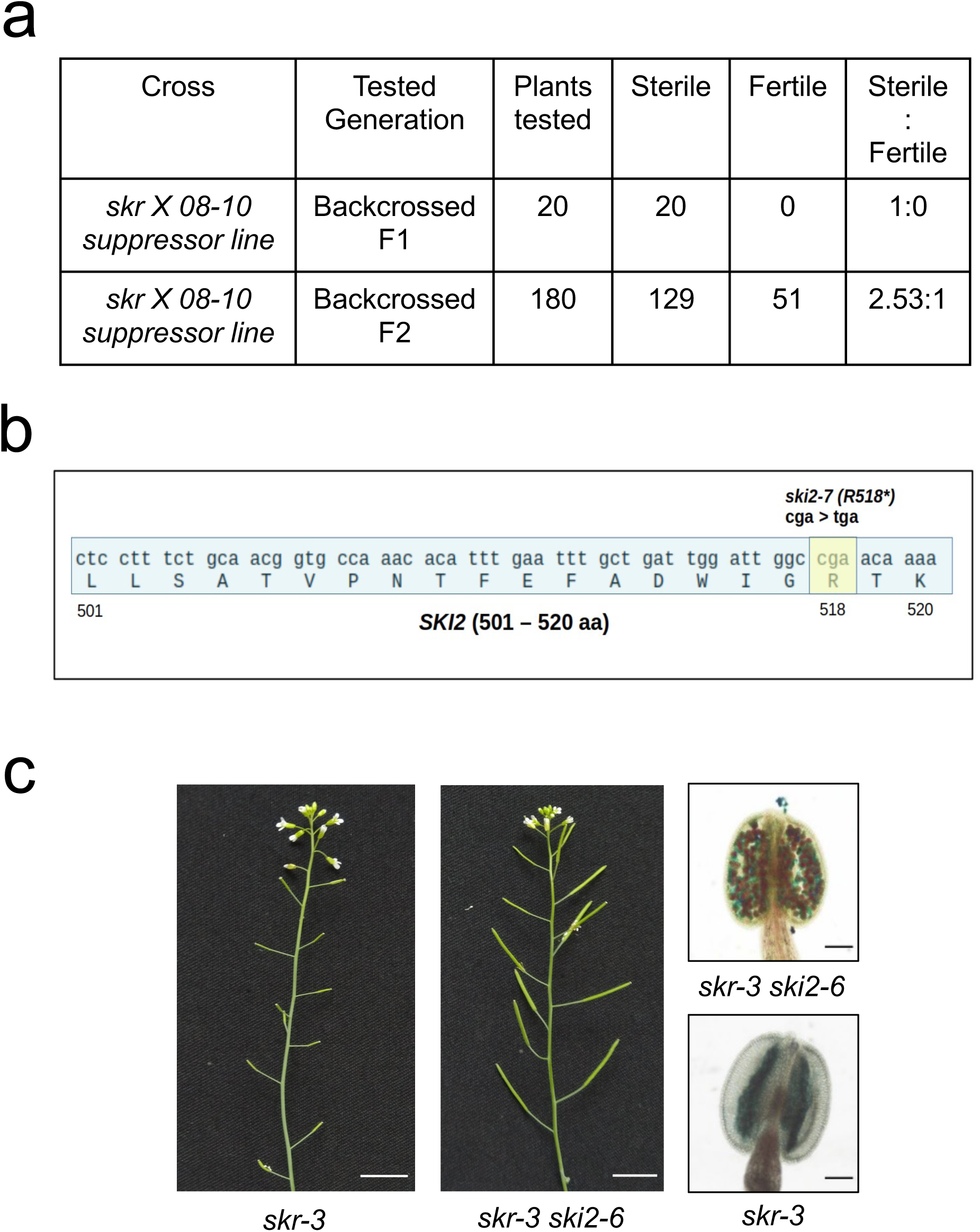
*ski2* is a suppressor of *skr*. (a) Mendelian segregation of the suppressor mutation in the 08-10 *skr* suppressor line (b) A C>T transition in the *ski2* allele (named *ski2-7*) present in the 08-10 *skr* suppressor line results in a TGA termination codon at position 518. (c) Restoration of fertility and pollen viability in *skr-3* by the independent allele *ski2-6*. Scale bar: inflorescence 1 cm; anthers 100 μm.

**Extended Data Fig. 7:**
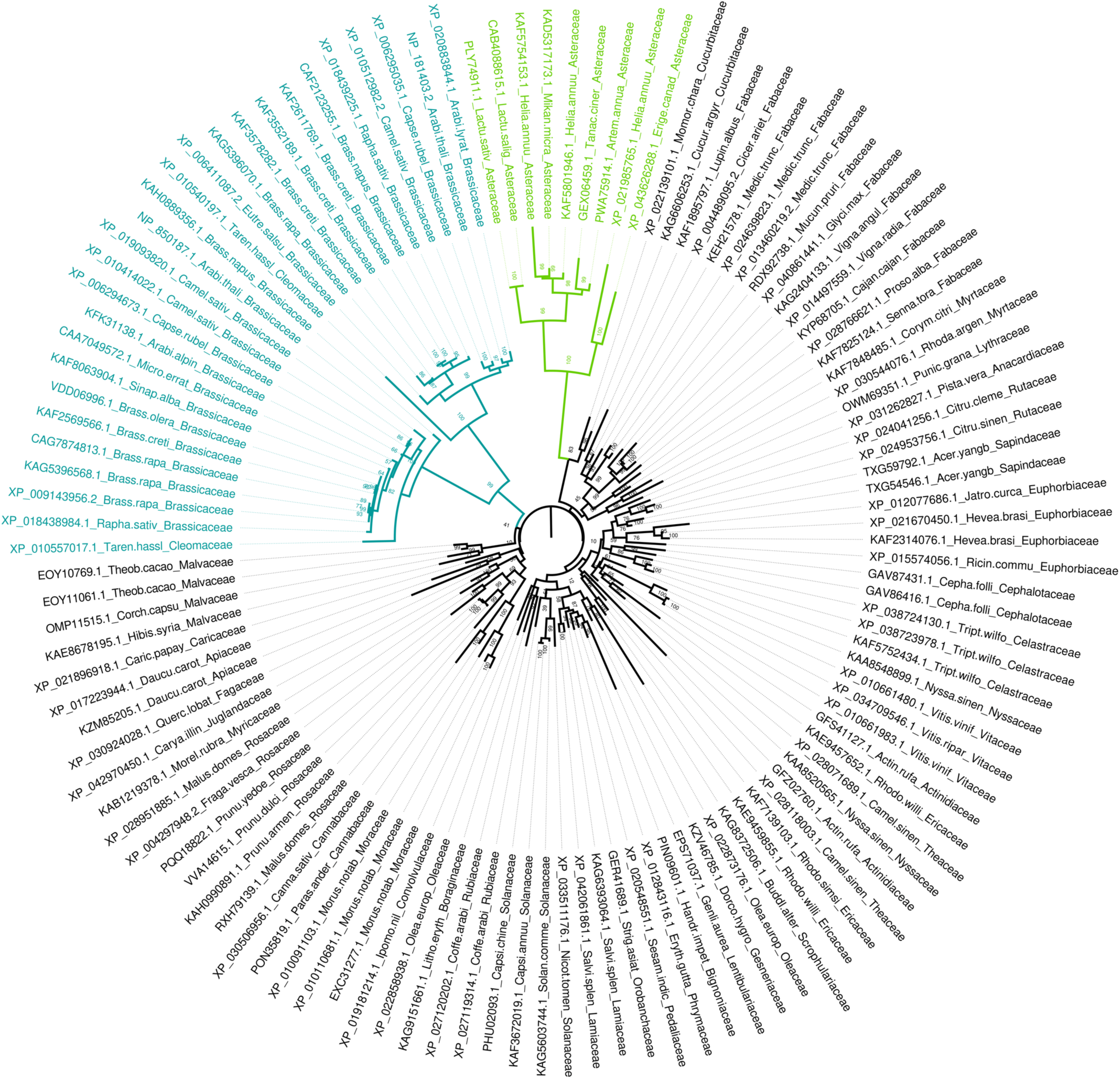
Phylogenetic tree of the SHUKR family from eudicots. 119 representatives from 85 species were used for tree construction using only the SHUKR domain that is shared by all representatives of the family. Genbank ID and species are indicated. Brassicaceae+Cleomaceae (dark green) and Asteraceae (light green)

**Extended Data Fig. 8:**
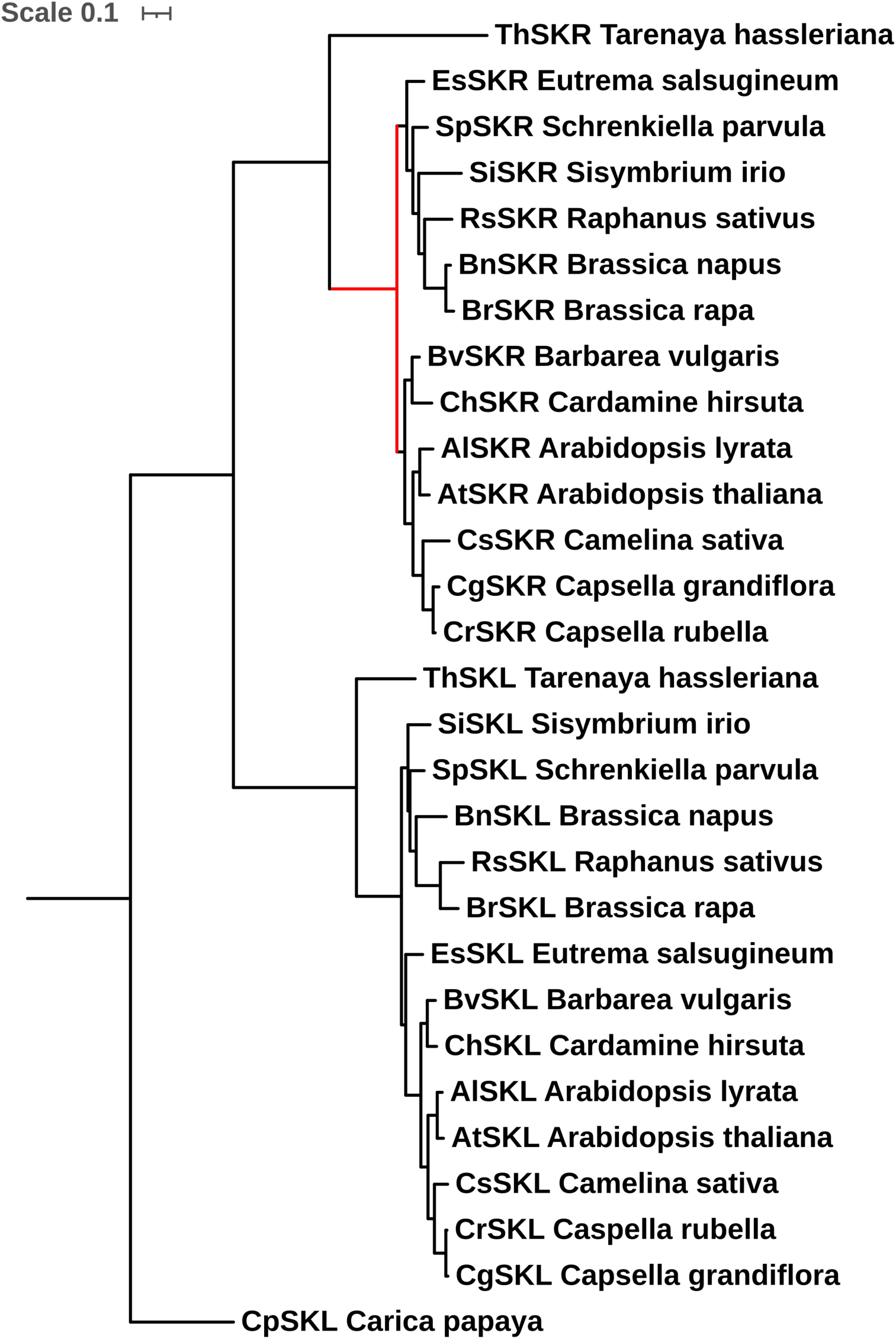
Phylogenetic tree of Brassicales *SHUKR* and *SHUKR-like* (*SKL*) genes used for branch-site test. The *SHUKR* gene family is represented by a single gene in *Carica papaya*, whereas Brassicaceae and Cleomaceae experienced a gene duplication event prior to their divergence. CDS were aligned using MAFFT, and RAxML was used to build the phylogenetic tree with 1000 bootstraps. The red branch was selected as foreground for *SKR*. Branch-site test for positive selection was performed using the CODEML package implemented in EASYCoDEML. Likelihood Ratio Test (LRT) results of Model A vs Model A null (Extended Data Table 1), gave a p-value of 0.0031 for *SKR*. The tree was unrooted.

**Extended Data Table 1:** Branch-site test results for positive selection in the SKR branch in the Brassicaceae. Branch site models postulate three classes of sites: those under purifying selection (0< ω<1), neutral (ω=1), and those under positive selection (ω>1) in the foreground branch being tested (Yang 2007)^53^. Model A is compared to Model A null in which the third class is set to ω=1. Positive sites are indicated together with posterior probabilities. P-value for likelihood ratio test (LRT) results are calculated from the χ^2^ distribution taking as critical value χ^2^= 2(LnL_MA_-LnL_MAnull_) and dividing by 2 (Yang 2007)^53^.

| Model | LnL | Estimates of parameters |  |  |  |  | Model compared | LRT P-value | Positive sites |
| --- | --- | --- | --- | --- | --- | --- | --- | --- | --- |
| Model A | -4394.302739 | Site class | 0 | 1 | 2a | 2b | Model A vs. Model A null | 0.003120857 | 7 E 0.932,22 F 0.672,25 V 0.529,31 E 0.613,43 F 0.553,57 E 0.962*,63 L 0.974*,82 K 0.594,100 P 0.747,112 R 0.668,114 C 0.643 |
|  |  | f | 0.66753 | 0.21979 | 0.08477 | 0.02791 |  |  |  |
| | | $\omega_0$ | 0.20115 | 1.00000 | 0.20115 | 1.00000 | | | |
| | | $\omega_1$ | 0.20115 | 1.00000 | 22.04175 | 22.04175 | | | |
| Model A null | -4398.670454 | 1 |  |  |  |  |  |  | Not Allowed |

## Methods

### Plant Material and Growth Conditions

Arabidopsis mutant strains used in the study were *skr-1* (GT_5_101780), *skr-2* (CSHL_GT5061) and *skr-3* (SALKSeq_034497.1), and *ski2-6* (SALK_122393)^91^. *skr-1* and *skr-2* alleles are in Ler ecotype and have a Ds insertion in AT2G38690; *skr-3* is in Col-0 ecotype. Plants were grown at 22 °C under 16 hours of light and 8 hours of dark in a 1:1:1 proportion of peat, perlite, and vermiculite soaked with 0.5x Murashige Skoog (MS) solution initially followed by irrigation during growth with water. For growth on plates and selection of transformants, seeds were surface sterilized and plated on MS plates supplemented with 2% sucrose and appropriate antibiotics.

### Coexpression screen for meiotic genes

We selected 23 known meiotic genes (DUET/MMD1, MS5/TDM1, DMC1, TAM, SDS, AML5, RAD51, RAD50, ASY1, ZYP1, REC8, SPO11-1, SPO11-2, PHS1, MRE11, MND1, OSD1, NBS1, MSH4, MSH5, HOP2, MER3, PRD1), and each of these genes was used as a query in the Expression Angler program^92^ to identify co-expressed genes. For each query, the top 100 hits were selected from each of three data sets, NASCArrays, Bio-analytic resource, and AtGenExpress Plus-Extended tissue compendium. We selected those genes common to three data sets and which appeared at least twice in independent queries. Single copy genes were selected using the Arabidopsis single copy genes list from the University of California, Davis (http://www.cgpdb.ucdavis.edu/COS_Arabidopsis/). We manually selected 29 genes that showed floral stage 9 peak/specific expression corresponding to meiosis stages from AtGenExpress Visualization Tool^93^. We tested one to two insertion alleles for each of the selected genes and for each of these alleles about 20 plants were genotyped by PCR to identify homozygous mutant plants and these were phenotyped by examining silique length and pollen viability by Alexander stain^94^.

### Plasmid constructs and transgenics

For complementation, a genomic fragment of 2220 bp (gSKR) comprising the *SKR* gene including 3’UTR was amplified from Ler ecotype and cloned into a modified binary vector pZP222 by Gateway cloning (Invitrogen). The binary construct was introduced into Agrobacterium strain AGL1 and transformed into *skr-1/+* and *skr-3/+* plants by in planta transformation^95^. *skr* mutant T1 plants were genotyped by PCR and tested for complementation. Plant vectors were ordered from (https://gatewayvectors.vib.be/index.php/gateway-vectors). The pro*SKR*:*GFP-gSKR* gene fusion construct was made by assembling the SKR promoter, GFP, and SKR genomic coding region plus 3’UTR using three-way Gateway technology in the binary destination vector pB7m34GW,0^96^. The pro*SKR:nlsGFP-GUS* construct comprised 1170 bp genomic DNA from 690 bp upstream of ATG upto 30 bp of the second exon fused in frame with GFP-GUS in the pBGWFS7 vector. F-box gene nlsGFP fusions were assembled in the pBnRGW_Redseed vector and introduced into plants. Single locus insert transgenic skr/+ lines were made homozygous for the transgene and examined in T4 for GFP fluorescence. The hairpin RNAi construct for PBE1^80^ was expressed under control of SKR regulatory elements comprising promoter and 3’UTR and contained the PDK intron from pBm42GWIWG8_1. The construct and a noRNAi control were introduced into *skr/+* plants and transgenics that were *skr* mutant were analysed for pollen viability. The *skr-1* mutant allele was used for all studies unless indicated otherwise. proMSP1:SKRcds constructs were made via overlap extension PCR, cloned into pENTR D TOPO vector using NotI and AscI restriction mediated cloning, and further cloned into pZP222 via Gateway cloning.

### Cytology and imaging

For Alexander staining, anthers from unopened buds were dissected out on a slide, placed under a coverslip, and Alexander stain was added from the side to fill the entire coverslip followed by sealing using nail-polish. The slides were incubated at room temperature for 3-4 days and visualized under 10x /20 x objective.Whole anther and ovule clearing was performed as in^97^, anther staging of plastic sections was performed as in^98^ and ovule staging as in^99^. Immunostaining for GFP:SKR in meiosis was performed as in^100^. Acid chromosomal spreads were prepared as described in^101^. GFP / DAPI fluorescence was visualized after mounting of meiotic anthers under the coverslip with 30-50 % glycerol / Antifade. ImageJ and Photoshop were used to edit and assemble images. Electron microscopy sample preparation and imaging was done as in^102^ and stripe artifacts in the EM images were removed using the method mentioned in^103^.

### Suppressor screening and analysis

*skr-1* mutant seeds (M0) were bulked from multiple plants and 0.3 % EMS solution was used to mutagenise the M0 seeds for 5 hours followed by extensive washing with water. Seeds from 934 M1 plants were collected individually and M2 families were screened for fertility restoration. One suppressor line 08-10 was identified and backcrossed to *skr* mutants to get BC1 F1 seeds. Seeds from BC1 F1 plants were collected and bulk segregant analysis on BC1 F2 plants was performed on fertile and sterile pools. Genomic DNA was isolated from the fertile and sterile pools (50 plants each) and used for whole genome sequencing.

To map the suppressor, the parental un-mutagenised *skr* genome was sequenced and *skr* specific reference genome was made by replacing the SNPs of *skr* to Ler-0 genome^104^ sequence using the *bedtools – consensus*^105^. Variants were called for the fertile and sterile bulk reads from the *skr* reference genome using *freebayes*^106^ and indels were removed. SNPindex was calculated using the R package QTLSeqr^107^ for the fertile bulk and sterile bulk. SNPs with a deltaSNP of fertile to sterile score greater than 0.5 were analysed for genic perturbations. The 08-10 line had an in frame stop codon in the SKI2 gene, at amino acid position 518. The suppressor phenotype was confirmed by generating the double mutant using the independent alleles *ski2-6* and *skr-3.* The *ski2* allele in line 08-10 was named *ski2-7*.

### Expression profiling

RNA from meiotic stage anthers was isolated using Qiagen kits for *skr-1,* Ler and *skr-1 ski2-7*. RNA sequencing was performed on Illumina platforms following the manufacturer’s instructions. Differentially expressed genes were found using DESeq2 keeping a FDR adjusted *P* value < 0.05^108^. Validation for eight upregulated F-box genes was performed by RT-qPCR (Supplementary Figure 3) which confirmed the RNAseq results. The tissue-specific expression analysis of deregulated genes was performed by downloading TPM values for each gene from CoNekT (https://evorepro.sbs.ntu.edu.sg) database^65^. Meiocyte RNAseq TPM value^109^ was combined with the CoNekT dataset. The tissue enrichment score was calculated by row normalization using the largest TPM score. Enrichment score of 1, in a tissue/cell type was taken to represent maximal expression in that type. The number of non-overlapping genes with enrichment score 1, was calculated and plotted for meiocytes, UNM, BCP, TCP/MP and others.

### Proteasome inhibitor treatment

MG132 and MG115 stocks at 20 mM concentration were made in DMSO. Inhibitor solutions were used at 100 µM concentration^110^ by dissolving in half strength MS where 0.5 % DMSO was present. Control solution was 0.5 % DMSO in half strength MS. Sections of freshly cut *skr* inflorescence stems were placed in 200 µl inhibitor or control solution in Eppendorf tubes for 3 hrs, followed by 60 hrs of recovery in half strength MS solution. Anthers were dissected and viable pollens were identified by Alexander staining. At least 100 anthers were analysed per sample (treatment and control) per experiment. Number of anthers with 10 or more viable pollen were counted per 100 anthers for treatment and control and the ratio was calculated. The experiment was conducted blind and repeated thrice. The ratio paired t test (two-tailed) on the logarithm of ratios was used to calculate p values (GraphPad).

### Sequence analysis

Sequence searches to detect homologs of the SHUKR domain were conducted using the PSI-BLAST (RRID:SCR_001010)^111^ and JACKHMMER (RRID:SCR_005305)^112^ programs with profile-inclusion threshold of expect (e)-value at 0.01 against the non-redundant database of National Center for Biotechnology Information (NCBI) clustered down to 50%. Clustering of proteins based on bit score density and length of aligned sequence was performed using the MMseqs program. Remote homology searches were performed using profile-profile comparisons with HHpred program (RRID:SCR_010276)^113^ against profile libraries from the PFAM (RRID: SCR_004726)^114^ and PDB (RRID:SCR_012820)^115^ databases. Multiple sequence alignments were built using Mafft program with maxiterate parameter set to 3000 followed by manual adjustments based on secondary structure prediction. Secondary structures were predicted using the JPred (RRID:SCR_016504)^116^ and RoseTTaFold^117^ programs. The globular domains in the protein were objectively identified using Shannon entropy-based compositional analysis with the SEG program with a window size of 45, low-cut of 3.4, high-cut of 3.75, and minimal length of the high-complexity segment of 15 residues.

The analysis of the Shannon entropy (H_j_) for a given multiple sequence alignment was performed using the equation:

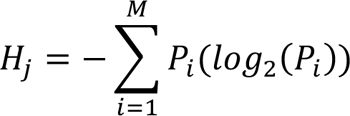

P is the fraction of residues of amino acid type i and M is the number of amino acid types. The Shannon entropy for the jth position in the alignment can range from 0 (only one residue at that position) to 4.32 (all 20 residues equally represented at that position). The reduced alphabet was created by the following grouping: h= L, I, M, V, A; a= F, Y, W; b=K, R, H; n=E, D; p=Q, N; o=T, C, S; t=P, G, d=-(gap).

### Phylogenetic analysis

SKR and SKL coding sequences of different species were extracted from Genbank, Phytozome, and CoGeBLAST^118,119^ (Supplementary File 1). Genomic fragments were annotated to CDS by the online tool MEGANTE^120^. For the branch site analysis, MAFFT was used to build the alignment and RAxML was used to build a phylogenetic tree with 1000 bootstraps^121,122^. Branch site model tests of the CODEML package^82^ were performed as implemented in EasyCoDEML^123^ by selecting the branch of choice. The tree was unrooted. Protein phylogenetic analysis was performed using the maximum likelihood method implemented in the IQtree program (RRID: SCR_017254)^124^ under multiple parameter regimes using: 1) the Q.plant substitution matrix derived from alignments specific to plant nuclear genes and 1 invariant site category with 8 gamma distributed sites; 2) the LG substitution matrix with 1 invariant site category with 8 gamma distributed sites or 3) with a 20-profile mixture model. Bootstrap values were calculated using the Shimodaira-Hasegawa-like approximate likelihood ratio (SH-aLRT) ^125^.

F-box gene sequences of 710 Arabidopsis genes used for ortholog analysis were obtained from Ref. 84 of which 64/69 were common to *skr* upregulated. Classification of LTS and STS genes was according to Ref. 84. Identification of orthologs was done using Proteinortho^126^.

## Acknowledgments

We thank David Twell for kindly providing the LAT52:GFP reporter line, ABRC and CSHL for seeds. Kathakali Bannerjee and Manoj Kamble for technical support, Mekala Sai Kiran for maintenance of plants, the NGS and Advanced Microscopy facilities at CCMB, and Priyanka Pant for help in illustrations. We are grateful to Elliot Meyerowitz, Utpal Nath, and Manjula Reddy for valuable comments on the manuscript.

## Author contributions

PS, SP, JND, JD and IS conceived and planned experiments. SP, PS, AR, JND, VS, HA, AS, and IS performed experiments. SP, PS, AR, AS, CK, HG, JD, and IS provided analysis of the experimental results. LA contributed the evolutionary analysis of Figure 6. IS and PS wrote the manuscript with contributions from LA and JD.

## Additional information

Supplementary information is available for this paper.

## Materials and correspondence

Correspondence and requests for materials to be addressed to IS. Sequencing data will be deposited at the European Nucleotide Archive.

## Supplementary information

**Supplementary Table 1:** Differentially expressed gene lists.

**Supplementary Table 2:** Male gametogenesis enrichment analysis of *skr* upregulated genes.

**Supplementary Table 3:** Analysis of genes differentially expressed in *skr*.

**Supplementary Table 4:** Differentially expressed translation-related genes in *skr ski2*.

**Supplementary Table 5:** Enrichment for short taxonomic scale (STS) F-box genes upregulated in *skr*.

**Supplementary Table 6:** List of primers

**Supplementary** Figure 1: Multiple sequence alignment of the SHUKR domain.

**Supplementary** Figure 2: Multiple sequence alignment of SKR protein sequences used for branch-site analysis.

**Supplementary** Figure 3: Validation for eight *skr* upregulated F-box genes by RT-qPCR.

**Supplementary File 1:** Sequences of *SHUKR* family genes used in branch-site analysis.

