## Supplementary figures and images for "Sporophyte Directed Gametogenesis via the Ubiquitin Proteasome System"

### Supplementary Figure 1

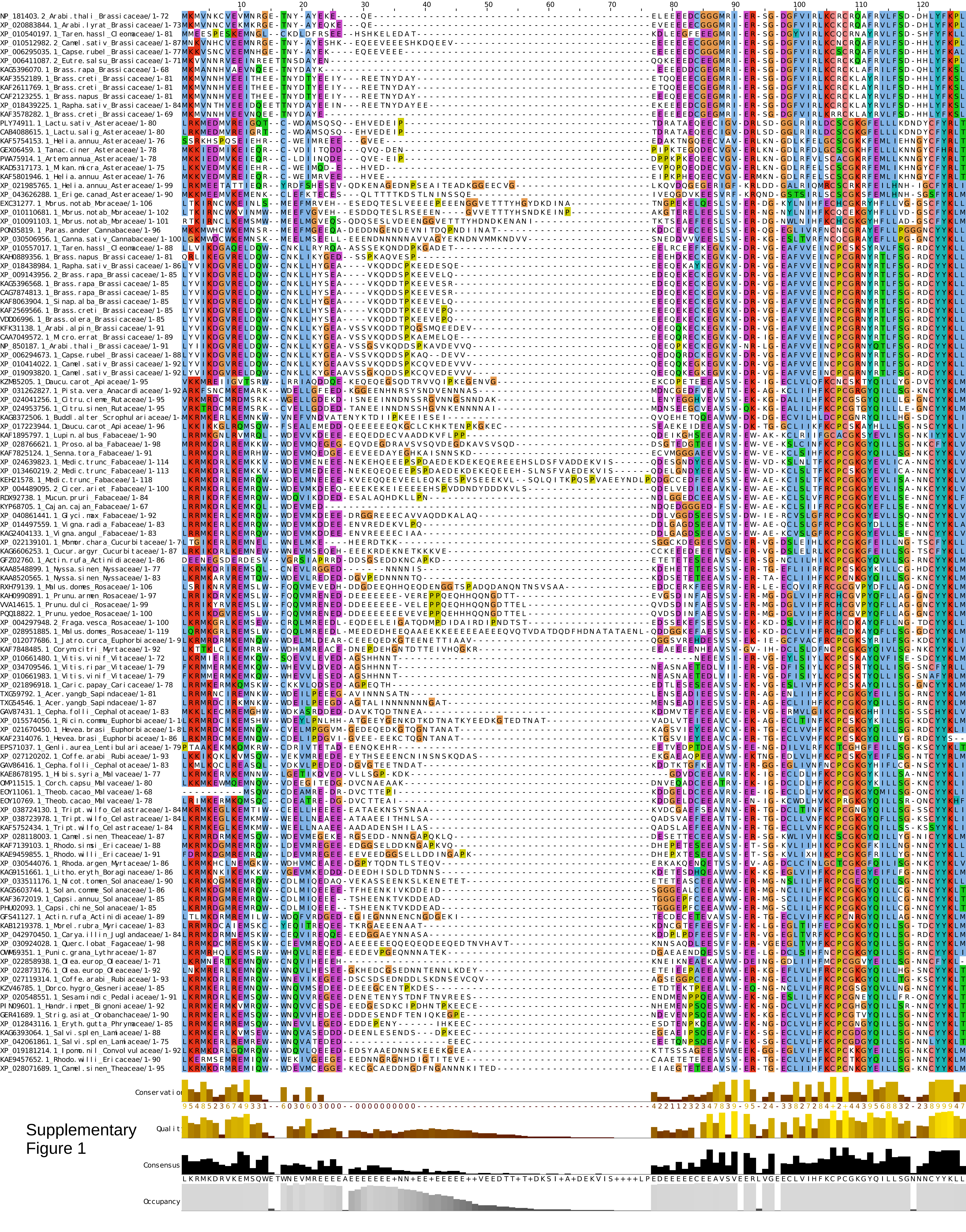


Supplementary Figure 1

### Supplementary Figure 2

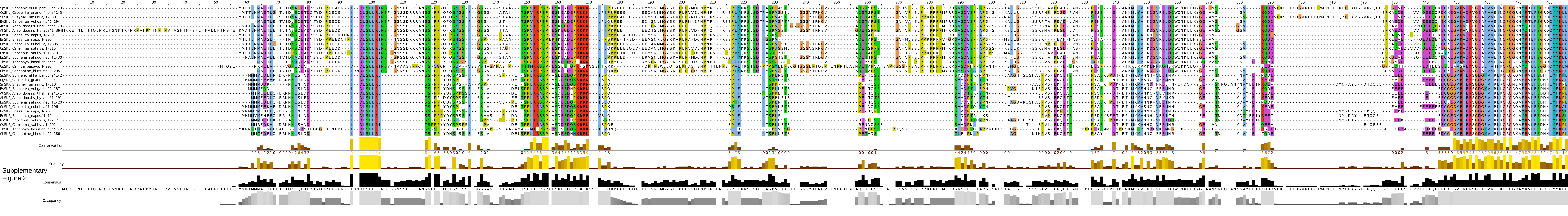


Supplementary Figure 2
