## Supplementary Figure 3 for "Sporophyte Directed Gametogenesis via the Ubiquitin Proteasome System"

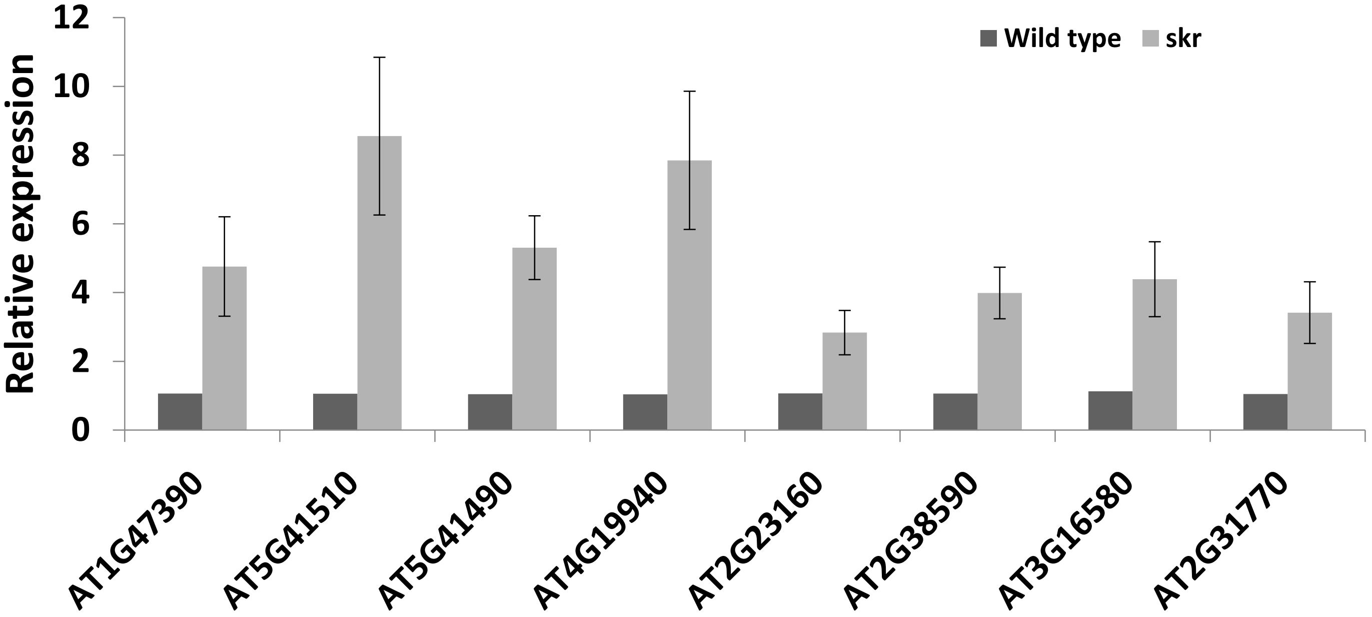


**Supplementary Figure 3:**
**Relative expression by RT-QPCR of 8 F-box genes upregulated in *skr* RNAseq**
Relative expression was calculated by the $2^{-DDCt}$ method (Livak and Schmittgen, 2001). Bars: SEM of 3 biological samples.
