## Supplementary File1 for "Sporophyte Directed Gametogenesis via the Ubiquitin Proteasome System"

### Supplementary File 1

Sequences used for Branch site analysis
>SpSKL Schrenkiella parvula
ATGACTTTAACTCTAAGCATGGCGGAAACCCTAGATACTTTGATCGATAACGCCGGAGAA
ACGGATTACTCCTACGATCATCCCGAGGAAGCAGATAACATTGATCTCTCTCTCCTTCGT
ATCAACAGCTTCGGTAACTCCTTAGATCGCCGCCGTGCGAACTCGTCTCCTCCGCAGTTT
AAATCGTACGGATCTTTCGGCTCTTCTTCCACCGCTGCCACAAGCCCCGTGAAACGCCCA
TCCCCGGAGTCGAAAGAAGCCGACGAGCCCAGACGCAAGAAGCTGTTTCTTCCACAATCG
GAGGAAGAAGAAGACGAAATGATGTCTAATCGCATGGGCTATTCGAAGATTCCTCTTCCT
ATGGATTGCAATCCGATGCGAATCCGCTCGTCTCCTATCTACAAGCGATCTCTCTCCGAC
TCATTTGCTTCGCCCGGTGCATCTACGTTCGGCGTCGCTCAAGGAACATCACCATCAGGT
AACACCGTCCCTTCTCTACCGCCACGGCCACCGGTATTCCGGAGGTCTGTCTCCGATGTA
TCGCCGTCACCTTCGAAGGCTCTTCTTGGATCTTCTCACTCCACTGCAGTTCCTGAAGCT
AAACTTGCGAATCCAGAAACCTCTGAAGCAAATAAGATGCTCTTTGTTATCAAGGACGGA
GTTCGTGAATTGGATCAGTGGTGTAACAAGCTCCTACAGTACGGTGAAGGAGTTTCTTCT
GTTAAACAAGATGACAGTCCTAAGGATCTTATCAAGGATGGAGTTCGTGAATTGGATCCG
TGGTGTAACAAGCTCATAAAGCACGGTGAAGCAGACTCTTTAGTCAAACAAGATGACAGT
CCTAAGGGAGAAGTTGAGTCACAAGAGGAGCAAGAGAAAGAGTGTAAAGAAGGCGTGAAG
GTGGACAGAGTCGGTGAGGCGTTTGTGGTTGAGATCAACTGTCCATGTGGAAGAAACTAC
CGAACTCTCTTCTCAGGCCGTGACTGCTACTACAAGCTCTTG
>CgSKL Capsella grandiflora
ATGACGACACTAAACATGGCGGAAACTCTAGGAACCTTAATCGATATCTCCCAAGAAACG
GATTGTACCTACGATCCTGACGAAGATGAAATCGATCTCTCCCTGCTTCGTCTCAACAGC
TTCGGGAACTCCTCCGATCGCCGCCGTGCGAACTCGTCTCCTCCGCAGTTTCAATCGTAC
GGGTCTTTCGGCTCATCTTCAGCCACCGCGATAACCAGCCCCGTGAAACGTCCATCGCCG
GAATCGAAGGAGGCTGACGAGCCGAGACGCAAGAAACTGTTTATTCCACGACCTGAGGAA
GAAGAAGAAGACGCGAATATGATGGGATATTCTAAGATTCCGCTTCCGGTGGTTGAATTG
AACCCGAATCGAATCCGTTCGCCTCTTTACAAGCGATCTTTATCCGATACATTTGCTTCG
CCCGTTGGATCTATACTCGGGTCGGGTAATACACGAAACGGCGTCGCTCAAGAAACATCA
CCGTCTTCGGGTAACGTCCCTTCTCTCCCGCCACGTCCACCGATGTTCCGGAGGTCCGTC
TCTGATGTTTCTCCGGCACCTTCCTCCAAGGCTCTTCTCCGATCTTCTCGCTTCAATCCA
ATTCCTGATGGTGACTTTGTGAATCCAGAAAGCTCTGAAGCCAATAAGATGCTGTATGTT
ATCAAGGATGGAGTTCGTGAATTGGATCAGTGGTGTAACAAGCTCCTTAAGTACGGGGAA
GCAGTCTCCTCTGTGAAACAAGACGACAGTCCCAAGGCACAAGATGAGGTAGTACAAGAG
GAACAACAGAGAGACTGTAAAGAAGGTGTGAAAGTGGATAGAGTCGGCGAGGCGTTTGTG
GTTGAGATCAATTGTCCATGTGGAAGAAACTACCGAACTCTCTTCTCTGGACGTGACTGC
TACTACAAGCTCTTG
>SiSKL Sisymbrium irio
ATGACGCTTACTCTAAGCATGGCGGAAACCCTAGATTCTCTGATCGATAACGCTCAAGAA
ACAGATTACTCCGACGATCATCCCGAGGAAGACAACATTGACCTCTCTCTCCTTCGTATC
AACAGCTTCGGAAACTCCTCAGATCGCCGCCGTGCTAACTCCTCTCCTCCACAGTTTCAA
TCGTACGGATCTTTCGGCTCTTCGTCAACCGCCGCCACAAGCCCCGTGAAACGTCCGTCA
CCTGAGTCGAAAGAAGCCGACGAGCCTAGACGCAAGAAGCTGTTTCTTCCTCCACCTGAG
GCTGAAGAAGACGAAAAGATGTCTACTCTCATGGGATACTCGAAGGTTCCTCTTCCTAAT
GATAGCAACTTGACGCGAAGCCGTTCGTCTCCCATCTACAAGCGATCTCTCTCCGATACG

TTTGCTTCTCCCGTGGCGTCTACGTTCGGGTCGGGTTATACACGTGGCGGCGTCGCTCAA
GAAACAGCTCCATCGGGTAATGTCGTCCCTTCTCTGCCGCCACGTCCACGGCCACCTCTC
TTCCGTAGGTCCGTCTCCGATCTCTCACCGGCACCTTCGAATGCTCTTCTTGGATCTTCT
TCAGAGACCTCTGAAGCCAATAAGATGCTCTATGTTATCAAGGATGGAGTTCGTGAATTG
GATCAGTGGTGTAACAAGCTCTTACAGTACGGTGAAGCAGTTTATTCTTCCAAACAAGAT
GATAGTCCTAAGTCTCTTATCAAGGACGGAGTACGTGAGTTGGATCAGTGGTGTAACAAG
CTCATTCAGTACGGTGAAGCTGTCTCCTCTGTCAAACAAGACGACAGTCCCAAGGAAGAA
GTTGAGTCACTAGAGGAGCAAGAGAAAGGGTGTAAAGAAGGAGTGAAGGTGGACAGAGTC
GGCGAGGCGTTTGTGGTTGAGATCAACTGTCCATGTGGAAGAAACTACCGAACTCTCTTC
TCTGGCCGTGACTGCTACTACAAGCTCTTG
>BvSKL Barbarea vulgaris
ATGGCGGAAACCCTAGAAACCGTAATCGACTTATCCCAAGAAACTGATTACACGTACGAT
CCCGAGGAAGATGACATTGATCTGTCTCTCCTTCGTCTCAACAGCTTCGGAAGCAACTCC
GATCGCCGCCGTGCAAACTCGTCTCCTCCGCAATTTCAATCGTACGGATCTTTCGGCTCC
TCTTCAACCACCAGCCCCGTGAAACGTCCATCACCGGAATCGAAGGAAGCTGACGAGCCT
CGACGCAAGAAACTGTTTCTTCAACGACCTGAGGAAGAAGATTCGAATCTAATGGGATAT
TCGAAGATTCCGCTTCCTATGGAATTCAATCCGACGCGAATCCGTTCTTCTCCTCTTTAC
AGGCGATCCTTATCCGATACATTTGCTTCGCCCGTTGGATTTACATTCGGGTCGGGTTAT
ATACGAAACGGCGTCGCTCAAGAAACAGCACCATCTTCGAGTAATGTCGCTTCGCTTCCG
CCACGGCCACCGATGTTCCGGAGGTCCGTCTCCGATGTTTCTCCTGCACCTTCTAAGGCT
CTTCTTGGATCTTCTCGCTTTACTGCAATTCCTGAAGCTGATCTTGTGAATCCAGAAAAC
TCTGAAGCCAATAAGATGTTGTATGTTATCAAGGATGGAGTTCGTGAATTGGATCAATGG
TGTAACAAGCTTCTAAAGTACGGTGAAGCAGTCTCCTCTGTGAAACAAGACGACAGTCCT
AAGGCAGAAGATGAGGTAGTAGAAGAGGAGCAGCAGAAAGAGTGTAAAGAAGGCGTGAAG
GTGGACAGAGTTGGTGAGGCCTTTGTGGTTGAGATCAACTGTCCTTGTGGAAAAAACTAC
CGAACACTCTTCTCAGGCCGTGACTGCTACTACAAGCTCTTG
>AtSKL Arabidopsis thaliana
ATGGCGACGCTAAACATGGCGGAAACCCTAGAAACCCTAATCGCTAATTCCCAAGAAGTT
GATTACTCCTACGATCCCGAAGAAGATGACATTGATCTCTCCCTCCTCCGTCTCAACAGC
TTCGGAAACTCCTCTGATCGCCGCCGTGCGAATTCCTCTCCTCCGCAATTCAAATCGTAC
GGATCTTTCGGTTCCTCTTCAACCACCGCAACAACCAGTCCCGTGAAACGTCCATCGCCG
GAATCGAAGCAAGGTGACGAGCCGAGACGCAAGAAGCTGTTTATTCCGAGACCTGAGGAA
GAAGAAGACCCGAATCTCATGGGATATTCAAAGATTCCGCTTCCTGTGGTTGAATTCAAT
CCGACGCAAATCCGTTCTCCTCTTTACAAGCGATCTTTATCCGATACATTTGCTTCGCCC
GTTGGATCTACATTCGGGTCGGGTGGGTCGGGTTATACCCGTAACAGCGTCGCTCAAGAA
ACATCACCTCCTTCGGGTAATGTCCCTTCTCTCCCGCCACGTCCACGGATGTTCCGGAGG
TCCGTCTCTGATCTTTCGCCGGCACCGTCCTCCAAGTCTCTTCTTGGATCTTCTCGCTCC
AATGCAATTCCTGAAGGTGATCTCGCGAATCCAGAAAGCTCTGATGCCAATAAGATGCTG
TATATTATCAAGGATGGAGTTCGTGAATTGGATCAATGGTGTAACAAGCTTCTTAAGTAC
GGTGAAGCAGTCTCCTCTGGCTCTGTGAAACAAGACGACAGTCCTAAGGCAGTAGATGAG
GTAGTACAACAAGAGGAGCAACCAAAAGAGTGCAAAGAAGGTGTGAAGGTCAACAGACTC
GGTGAGGCCTTTGTGGTTGAGATCAACTGTCCATGTGGAAGAAACTACCAAACTCTCTTC
TCAGGCCGTGACTGCTACTACAAGCTCTTG
>AlSKL Arabidopsis lyrata

ATGAAACGTGAAATTAATTTGATAATTATCCAACTAAACCGGTTGTTTAGCAATAAAACC
CGGTTCAATCGACCCGCATTTCCCTATATAAACCCAACTCCAGTGATCGTCTCTTTCATC
AACTTCTCATTTCTCACTTTCGCATTGAACTTTATTAACAGTACTGAGATCAAGATGGCG
ACGCTAAACATGGCGGAAACCCTAGAAACCCTAATCGCTAGCTCCCAAGAAGCTGATTAC
TCCTACGATCCCGAAGAAGATGACATTGATCTCTCCCTCCTTCGTCTGAACAGCTTCGGA
AACTCCTCTGATCGCCGCCGTGCGAATTCCTCTCCTCCGCAATTCAAATCGTACGGATCT
TTCGGCTCCTCTTCAACCACCGCAACAACCAGCCCCGTGAAACGTCCATCGCCGGAATCG
AAGGAAGGTGACGAGCCGAGACGCAAGAAACTGTTTATTCCACGACCTGAGGAAGAAGAA
GACACAAGTCTCATGGGATATTCAAAGATTCCACTTCCTTTGGTTGATTTCAATCCGACG
CAAATCCGTTCTCCTCTTTACAAGCGATCTTTATCGGATACATTTGCTTCGCCCGTTGGA
TCTACATTCGGGTCGGGTTATACCAGAAACAGCGTCGGTCAAGAAACATCACCGCCTTCG
GGTAATGTCCCTTCTCTCCCACCACGTCCACCAATGTTCCGGAGGTCCGTCTCCGATCTT
TCGCCGGCACCTTCCTCCAAGTCTCTTCTTGGATCTTCTCGCTCCAATGCAACTCCTGAA
GGTGATCTTGTGACTCCAGAAAGCTCTGATGCCAATAAGATGCTGTATATTATCAAGGAT
GGAGTTCGTGAATTGGATCAATGGTGTAACAAGCTTCTTAAGTACGGTGAAGCAGTCTCC
TCTGGCTCTGTGAAACAAGACGACAGTCCTAAGGCAGAAGATGAGGTAGTACAAGAGGAG
CAACCGAAAGAGTGCAAAGAAGGGGTGAAGGTGAACAGACTCGGTGAGGCCTTTGTGGTT
GAGATCAACTGTCCATGTGGAAGAAACTACCAAACTCTCTTCTCAGGCCGTGACTGCTAC
TACAAGCTCTTG
>BnSKL Brassica napus
ATGACTCTAACCCTAAGCATGGCGGAAACCCTAGATGCTTTAATCGATAACTCCCAAGAA
ACCGATTACTCCTCCGCTCACCCCGAGGAAGACAACACTCAGAACATCGATCTCTCTCTC
CTCCGTATCAACAGCTTCGGTAACTCCTCCGATCGCCGCCGCGCGAACTCGTCTCCTCCG
CAGTTTCCATCGCACGGATCTTTCGGGTCTTCCTCCCCCGCCGCCGCCACCAGCCCCGTG
AAACGCCCCTCCCCGGAGTCGAAAGAAGCCGACGAGCCTAGACGCAAGAAGCTGTTTCTT
CCCCCACCACGTGACGCCGAGGATGACGAAACGATGTCGAATCGCTTGGGATACTCGAAG
ATTCCGCTTCCTGTGGATTCTAATCCGACGCGAGTCCGTTCGTCTCCGATCTACAGACGA
TCTCTCTCCGATACGTTTTCTTCCCGAAGCGGCGTCGCTCAGGAAACAGCACCGTCTCTC
CCGCCACGGCCACCGGTCTTCCGGCGGTGCGTCTCCGATGTTTCGCCGGCGTTTCATGGA
ACTGATCTTGCGAATCAAGAAACGTCCGAAGCCAATAAGATGCTCTATGCTATCAAGAAT
GGAGTTGTTGAATTGGATCAGTGGTGTAACAAGCTTCTACAGTACGGTGAATCGGTGAAA
CAAGATGGCAGTAGTCCTAAGGCACAAGTTGAGTCACCAGAGGAGGAGCACGACAAAGAG
TGTAAAGAAGGAGTGAAGGTGGACAGAGTTGGTGAGGCTTTTGTTGTTGAGATCAACTGT
CCTTGTGGAAGAAACTACAGAACTCTCTTCTCTGGCCGTGACTGCTACTACAAGCTCTTG
>BrSKL Brassica rapa
ATGACTCTAACTCTAACCATGGCGGATTCTCTGATCAACACCTCCCAAGAAACTGATTAC
ACCTACGATCATCCCCCCGAGGAAGATAACACTCCCATCGACCTATCTCTCCTCCGTATC
AACAGCTTCAACAACTCCTCAGATCGCCGCCGCGCGAACTCCTCTCCTCCGCAGTTCCCA
TCTCACGGATCTTTCTCCTCTCCCCCCGCCGCCGCCACAAGCCCCGTGAAACGCCCATCC
CCAGAGCCGACGAAAGAAGCCGACGAGCCGAGGCGCAAGAAGCTGTTTCCTTCCCCGCC
G
GAGACTAAGGAAGACGAGGAGATGTCTAACCGCCTGGGCTACAAGAAGGTCCCGCTTCCG
GCGGATTGCAATCCGGCGCGAATCCGTTCGTCTCCTGTCTACAAGCGATCTCTCTCCGAT
ACATTTGGATCGCGAGTCGCTCAAGAAACAGCACCATCGGGGAACATGGTCTCTTCTCTG

CCGCCGCGGCCACGGAGACCGGTGTTTGGGAGGTCCGTCTCCGATCTGTCGCCGGCACC
T
TCGATGTCGCTTCTTGGATCTTCTTCTCGTGAAGCTAATCATGCGAGTCAAGAAACAGCT
GAAGCCAATAAGATGCTGTATGTGATCAAGGATGGAGTTCGTGAGTTGGATCAGTGGTGT
AACAAGCTCTTACACTACAGTGAAGCAGTGAAACAAGATGATAGTCCTAAGGAAGAAGTT
GAGTTGCAAGAGGATGAGCAAGAGAAAGAGTGTAAAGAAGGGGTGAAGGTGGACAGAGTA
GGTGAGGCGTTTGTGGTTGAGATCAACTGTCCATGTGGAAGAAACTACCGAACTCTCTTC
TCTGGCCGTGACTGCTACTACAAGCTCTTG
>CrSKL Caspella rubella
ATGACGACACTAAACATGGCGGAAACTCTAGGAACCTTAATCGATAACTCCCAAGAAACG
GATTGTACCTACGATCCTGACGAAGATGAAATCGATCTCTCCCTCCTTCGTCTCAACAGC
TTCGGGAACTCCTCCGATCGCCGCCGTGCGAACTCGTCTCCTCCGCAGTTTCAATCGTAC
GGGTCTTTCGGCTCATCTTCAGCCACCGCGATAACCAGCCCCGTGAAACGTCCATCGCCG
GAATCGAAGGAGGCTGACGAGCCGAGACGCAAGAAACTGTTTATTCCACGACCTGAGGAA
GAAGAAGAAGACGCGAATATGATGGGATATTCTAAGATTCCGCTTCCGGTAGTTGAACTC
AACCCGAATCGAATCCGTTCGCCTCTTTACAAGCGATCTTTATCCGATACATTTGCTTCG
CCCGTTGGATCTATACTCGGGTCGGGTAATACACGAAACGGCGTCGCTCAAGAAACATCA
CCGTCTTCGGGTAACGTCCCTTCTCTCCCGCCACGTCCACCGATGTTCCGGAGGTCCGTC
TCTGATGTTTCGCCGGCACCTTCCTCCAAGGCTCTTCTCCGATCTTCTCGCTTCAATCCA
ATTCCTGATGGTGACTTTGTCAATCCAGAAAGCTCTGAAGCCAATAAGATGCTGTATGTT
ATCAAGGATGGAGTTCGTGAATTGGATCAGTGGTGTAACAAGCTCCTTAAGTACGGGGAA
GCAGTCTCCTCTGTGAAACAAGACGACAGTCCCAAGGCACAAGATGAGGTAGTACAAGAG
GACCAACAGAGAGACTGTAAAGAAGGTGTGAAAGTGGATAGAGTCGGCGAGGCGTTTGTG
GTTGAGATCAACTGTCCATGTGGAAGAAACTACCGAACTCTCTTCTCAGGACGTGACTGC
TACTACAAGCTCTTG
>CsSKL Camelina sativa
ATGACGACGCTAAACATGGCGGAAACCCTAGAAACCCTAATCGATAACTCCCAAGAAACG
GATTACACGTACGATCCCGAAGAAGATGACATTGATCTCTCACTCCTCCGTCTCAACAGC
TTCGGGAACTCCTCCGATCGCCGCCGTGCGAACTCGTCTCCTCCGCAGTTTCGATCATAC
GGATCTTTCGGCTCTTCTTCGACCGCCGGAATAACCAGCCCCGTGAAACGTCCATCGCCG
GAATCGAAGGATGCTGAAGAGCCGAGACGCAAGAAGCTGTTTCTTCAACGGCCTGAGGAG
GAAGAGCAAGAAGTAGAAGAAGACGCGAATCTCATGGGATATTCTAAGATTCCGCTTCCG
GTGGTTGTTCTCAACCCGAATCGAATCCGTTCGCCTATTTACAAGCGATCTTTATCGGAT
ACATTTGCTTCGCCCGCTGGATCTATGCTCGGGTCGGGTAATTCACGAAACGGCGTCTCT
CAAGAAACATCGCCGCCTTCGGGTAATGTCCCTTCTTCTCTCCCGCCGCGTCCACCGATG
TTCAGGAGGTCCGTCTCCGATGTTTCTCCCGCACCTTCCTCTTCCAAGTCTCTTCTCCGA
TCTTCTCGCTCCAATGCAATTCGTGAAGGTGATTTTGCGAGTCCAGAAAGCTCTGAAGCC
AATAAGATGCTGTATGTTATCAAGGATGGAGTTCGTGAATTGGATCAATGGTGTAACAAG
CTCCTTAAGTACGGTGAAGCAGCAGTCTCCTCTGTGAAACAAGACGACAGTCCTAAGTGC
CAGGTAGAAGATGAGGTAGTAGTACAAGAGGAGCAACAGAAAGAGGGTAAAGAAGGTGTG
AAGGTGGATAGAGTCGGCGAGGCCTTTGTGGTTGAGATCAACTGTCCATGTGGAAGAAAC
TACCGAACTCTCTTCTCAGGCCGTGACTGCTACTACAAGCTCTTG
>RsSKL Raphanus sativus
ATGACTCTGACTCTAAACATGGCGGATACTCTAATCAACCCCTCCCAAGAAGCGGATTAC

ACCTACGATCATCATCCCGAGGAAGATAACACTCCCATCGACCTCTCCCTCCTCCGCATC
AACAGCTTCGGAAACAACTCCTCAGATCGCCGCCGCGAACTCCTCTCCTCCGCAGTTC
CCATCTCACGGATCTTTCTCCTCTTCCTCCACCGCCTTCGCCACCACGAGCCCCGTGAAA
CGCCCCTCACCGGAGCCGAAAGAAGCCGACGAGCCGAGGCGCAAGAAGCTGTTCCTCTC
C
CCGCCGGAGACTAAGGAAGAAGACGAAGAAGAAGAGATGTCTAACCGCCTGGGGTATAAG
AAGGTCCCGCTTCCTCCGGATTGCAACCCGACGCGAGTCCGTCCGACTCCGGTCTACCAG
AGATCTCTCTCCGATACGTTTGCTTCCACCGGTTATACTCGTCGCCGCGGCGTTGCTCAA
GAAACAGCACCGTCGTCGGGTAACATCGTCTCTTCTCTGCCGCCGCGGCCACGCAAACCG
GTGTTCCGGAGGTCGGTCTCCGATCTTTCGCCGGCACCACCTTCGATGGCGCTTCTCGGC
TCTTCTTCTCGCTCCACTGCGGTTGATGAAGCAATTTTCGCGAATCGAGAAACCTCTGAG
GCCAATAAGATGCTGTATGTGATCAAGGATGGAGTTCGTGAATTGGATCAGTGGTGTAAC
AAGCTCTTACACTACGGTGAAGCTGTGAAGCAAGACGACTGTCCTAAGGAAGAAGATGAG
TCGCAAGAGGAGGAGGAGCAAGAGAAAGCGTATAAAGAAGGCGTGAAAGTGGACAGAGTG
GGCGAGGCGTTTGTTGTTGAGATCAACTGTCCTTGTGGAAGAAACTACCGAACTCTCTTC
TCTGGCCGAGACTGCTACTACAAGCTCTTG
>EsSKL Eutrema salsugineum
ATGGCTCTAAACATGGCGGAAGCCCTAGAAACCGTAATCGATAACGCTCAAGAAACGGAT
TACTCGTACGAACCAGAGGAAGATGACATCGATCTCTCTCTCCTTCGTCTCAACAGCTTC
GGGAGGTCCTCCGATCGATGCCGCGCGAACTCGTCTCCTCCGCAGTTTCAATCGTACGGA
TCTTTCGGATCTTCTTCAACCGCTGCCACCAGCCCCGTGAAACGCCCATCGCCGGAGTCG
AAGGAAGCCGACGAGCCCAGGCGCAAGAAGCTGTTTCTTCCCCGACCTGAGGAAGACGAA
AGGATGTCTAATCTCATGGGCTATTCGAAGGTTCCGCTTCCCATGGATTGCAATCCGACG
CGAATCCGTTCGTCTCCTATCTACAAGCGATCTCTATCCGATACGTTTGCTTCTCCCGGT
ATATCCACATTCGGGTCGGGTTATACCCGAGACGGCGTCGCTCAAGAAACAACACCCTCG
GGTAATATCGTCCCTTCTCTTCCGCCACGGCCACCGATGTTTCGGAGGTCCGTCTCCGAT
GTTTCGCCGTCACCTTATAAGACTCTTGGATCTTCTCGCTCGACTGCAATTCCTGAAGCT
GATTTTTCGAATCCAGAAGCCTCTGAAGCTAACAAGATGCTGTATGTGATCAAAGACGGA
GTCCGTGAATTGGATCAATGGTGTAACAAGCTCATAAAGTACGGTGAAGCAGTCTCTTCT
GTCAAGCAAGATGACAGTCCTAAGGGAGAAGTTGAGTCACAAGAGGAGGAGCAGCAGAAA
GATTGTAAAGAAGGAGTTAAGGTGGACAGAGTCGGCGAGGCCTTTGTGGTTGAGATCAAC
TGTCCCTGTGGAAGAAACTACCGAACTCTCTTCTCAGGCCGTGACTGCTACTACAAGCTC
TTG
>ThSKL Tarenaya hassleriana
ATGGCTGAAACCCTAATCGAAAACCACCAAGAAGCGGACTACTCGTACGAGCTCGAGGAA
GAAGAGGACATCGATCTCTCACTCCTCCGTCTCAACAGTTTCGGCAGCTCCTCCTCCGAT
CGCCGTCGATCTAACTCTTCTCCTCCGAAGTTCCACTCGAGCGGCGGATCTCTCAGCTCC
TCCCCCTATGCCGCCGTATCCGTCGGAAGTCCCGGTAAACGCCCCTCTCCAGAGTCCAAA
GATTCCGACGGGCCTCTCCGCAAGAAACATTTTCTCCGGCCGGCGGAAGACGATGCGAAC
CCTAATCTCCAGGGTTATACGAAGATTTCGCTTCCTATTGATCTCAGCCCGAACCGGACC
CGTTCACCTCTTCTCTTCAGGCGATCCCTTTCGGATTCGTTTGCTTCGTCCGGTCATGCT
AGACCAGTCGCCCAAGAAGCGTCACCTTCGGGTAATGTCGCTCCTCCACTCCCTCCACGG
CCACCGTTGTTCCGGAGGTCCGTCTCGGATATCACTCCGGCGCCGGCGAAGACTTTTCGG
GGGTCTTCCAGCTCCGTTGCTATCCCAGAAGCTGATCTTGGGAAGATGGGAACCTCCGAA

GCTGATAAGATGTTGTTGGTTATAAAGGATGGAGCGCAGGAGTTGGATCAATGGTGTAAC
AAGCTCCTTCGGTACAGGCAAGCAGCTTCTTCCTCCGAGAAGCAAAACGATGATCCCAAG
GGGGCAGATGAAACGGAAGAACTGAGGTGTGAAGAGTTCAAGGAAGGAGTGAAGGTAGAC
AGGGTCGGGGAGGCTTTTGTGGTTGAAATCAACTGCCCTTGCAGCAAATCCTACCGCGTT
CTGTTCTCCGGCCGCGACTGCTACTACAAGCTCCTC
>CpSKL Carica papaya
ATGACTCAATATGAAAATAAGAATGAGTTGGTAGTAGGCAATCTTCAAGAAAATGTGGAT
TTATCCCTCCTCCGCATCAACAGCTATAATGCCGCCGCATCGTATCCTCCCTGCACCTCT
TGTGGCTGCAACTTCTCCTGCGCCGTCTCCTCAGTCTCCAACGCCGGCGCCCCAGCCTCC
ACCCCCACCACCACCATGAAGCGCTCATCCCCTGAATCCTCCCTTTCCACTCAACCCAAA
TCCAAGAACTCTTCTCACATCAAGAGACGCCAACCTATCCCCTCTTGGCTTCTCCAGATT
TCCCTTCCCTTCGCTTCATTCTCTAACCCCACCCACACCCTCAACCCCTCCACCCCCGTT
CTTCATCGTTCCATTTCAGATCCTTATCCCTCCTACCTCGATCCTTCATGTGGACACGGA
TCAGGTGAACCGACACAGTCACCTATAGAGAATCCAAGAATAGAAGCTAGCAATCAGGAG
ACAGCTCCCGTGGCCTTCTCTGCGCCAAAGGGAAGTGGGCCTTCTGCATTGCCACCACTG
CCGCCCTCTCTGAGGAGATCCGCGTCCGACCCGAACCCGTCGCCCGCTAACACATTTTCC
ATGGGAAAAGATTCCATCAAGGAGGAGAGTCCCGATACTAAGAGGCTGAGGAGAATGAAG
GAACGCATGAAGCAGATGAGTAAATGGTGCAAGAAAGTGCTGCAAGATAGTGAAGATGCT
GGTCCTGAACAGACACATGAGGATACTCTTGAATCAGAATCTGAAGAAGCTGTGAGTGTG
GAGAAAGTTGGAGAGAGTTTGATTGTTCACTTCAAATGTCCCTGCAGCAAAGCGTATCAA
ATCCTTCTCTCTGGAGGGAACTGTTATTACAAGCTCATG
>ChSKL Cardamine hirsuta
ATGGCGGAAATCCTAGAAACCGTAATCGATTTATCCCAAGAAACAGATTACACCTACGAT
CCCGAGGAAGATGACATTGATAATGATCTGTCTCTCCTTCGTCTCAATAGCTTCGGAAGC
AACTCCGATCGCCGCCGTGCGAACTCGTCTCCTCCACAGTTCCAATCGTACGGATCTTTC
GGCTCCTCCTCAACCACAAGCCCCGTGAAGCGTCCATCCCCGGAATCGAAGGAAGCTGAC
GAGCCGAGACGCAAGAAGCTGTTTCTTCAACGACCTGAGGAAGAAGATTCGAATCTCATG
GGATATTCGAAGATTCCGATTCCTGGGGATTTCAATCCGACGCGAATCCGCTCTTCTCCT
CTCTACAAGCGATCCTTATCCGATACATTTGCTTCGCCCGTTGGATTCACATTCGGGTCG
GGTTATACACGAAACGACGTCGCTCAAGAAACAGCACCTTCGTCGAGTAATGTTCCTTCT
CTTCCGCCACGGCCACCGATGTACCGGAGGTCCGTCTCCGATGTTTCTGCGGCACCTAAG
GCTCTTCTTGGATCTTCTCGCTCAACTGCAATTCCTGAAGCTGATCTTGTGAATCCAGAA
AGCTCTGAAGCCAATAAGATGCTGTATGTTATCAAGGACGGAGTTCGTGAATTGGATCAA
TGGTGTAACAAGCTCCTAAAGTACGGTGAAGCAGTTTCCTCTGTGAAACAAGACGATAGT
CATAAGGCAGAAGATGAGTTAGTACAAGAGGAGCAGCAGAAAGAGTGTAAAGAAGGCGTG
AAGGTGGACAGAGTTGGTGAGGCCTTTGTGGTTGAGATCACCTGTCCTTGTGGAAGAAAC
TACCGAACACTATTCTCAGGCCGGGACTGCTACTACAAGCTCTTG
>SpSKR Schrenkiella parvula
ATGATGATGGTAGAAGATGAATTTCGAGACCGTAACAATCTGAGCATCAACGACATTGAT
CTCTCTCTCCTCCGCCTCTCCTCCCCACCTTACAACTGCTCTCGCTCCTCCTTCTTCTCC
ACCGTTTCTCCTGAGATCAGCCCCTTGAAACGCTCCTCCCCTGTTTCCGAAGATTCCGAT
CAGCCCAAACGCAGGAGAATCTCCCCCAAAAACCCAATCTTTATCACCTCTCCTCTTCGC
TCCACTCACCCGGAAATCCAATCTTCCTCGAAAGACGATCCGACCCGGAACCCGAATCTC
GCCGCCGGCCACATCAGTTGTTCCCACGCAAGCCCTGTTTCAGAGAAGCAAGAAACGTCG

CCGTCTGCTTCCAAGAGTCCTGATACAGAGACGATGAAGGTGGTTAACAATCACGTGGAG
GAGATCAATCGCGAAGAAACTAACTTTAATGCTTATGAGAATCAGCAGGAGGAGGAAGAA
GAAGAAGATTGTGGAGAAGGGATGAGGATAGAGAGATCTGGAGATGGATATGTAATTAGA
TTGAAGTGTAGATGCAAACAAGCTTTTAGGGTTCTTTTCTCTGATCATCACCTCTACTTT
AAGCATCTC
>CgSKR Capsella grandiflora
ATGATGATGATGATGGAAGAAAGAGCCGAAGATCGTAACCACAACCTGACCCTCGACGAC
ATTGATCTCTCTCTCCTCCGTCTCACTTCTCCACCTTACGACTACTCACCCTTCTTATCC
GCCGATGAAATCACCCCTCCGTTGAAACGCGCCTCCCCTGTTTCCGACGATTCCGATCAG
TCCAAACGAAGAAAACTCTCCCCTAAAGACCCGATCTTTATCACCTCTACTGTTCTCTTC
ACTTCCCTGGAGAACCAATCTTCATCTGTCCATGACCCGACTCGGTACACGAACCTCACA
AGCCCTGTTTCAGAGAAGCAAGATACGACGCCGTATGCTTCTAATACAGAGACGATGAAG
AAGGTTAGCAATTGCGTAGAGGAGATGAATCATGGAGAGGCTAACTATGCTTATGAGAAG
GAGCAAGAGGAAGTGGGAGAAGAAGAAGAAGAAGATTGTGGAGGAGGGATGAGGATA
GAGAGATCTGGAGATGGGTTTGTAATTAGACTCAAGTGTAGTTGCAGACAAGCTTTTAGG
GTTCTTTTCTCTGATCATCACCTCTACTTCAAGGCTCTC
>SiSKR Sisymbrium irio
ATGATGATGATGCAAGACCGTAACAATCTGAACATCAACGACATTGATCTTTCTCTCCTC
CGCCTCTCCTCTCCACCTTACGACTACTCTCGCTCCGCCGTTTCTCCTGAGATCAGCCCT
TTGAAACGCTCCTCCCCTGTTTCCGAAGAATCCGATCAGTCCAAACGAAGGAGAATCTCC
CCTCAAAACCCAATCTTTATCACCTCTCCTCCTCCTCAGTCCTCCACCATCGACGACCCG
ACCCCGTACCCGAATCTCACAGCCGCAAGCCCTGTTTCGGAGAAGCATGAAACGTCACCG
TCTGCTTTCAAGACTCCTGATAAAGAGACGATGAAGATGGTCAACGATCACAACCGCGAA
GAGACTAACTGTGATGTTTATGAGGAGAGCAATCGCCAAGAGGCTAACTATAATGCTTAT
GAGGAGATCAATCGCCAAGAGGCTGACTATAATGCTTATGAGGATCATCAGCAAGAGGAG
AAGGAAGAAGAAGAAGAAGAAGAAGAAGATTGTGGAGAAGGGATGAGGATAGAGAGATCT
GGAGATGGATTTGTAATTAGATTAAAGTGTAGATGTAACCTAGCTTTTAGGGTTCTTTTC
TCTGATCATCACCTCTACTTCAAGCGTCTC
>BvSKR Barbarea vulgaris
ATGATGATGATGGAACAAGACAATCTGAGTCTCGACGACATTGATCTCTCTCTCCTTCGC
CTCACTTCCCCACCTTACGATCACTCATCATCCTTCTTCTCCGCCGATGAACTCAGCCCT
CCTTTGAAACGCTCCTCCCCTGTTTCCGACGAATCCGATCACCCCAAACGCAGAAAACTC
TCCCCTCAAAACCCAATTTTTATCACCTCTCCTCTTCTCTTCACTTCCCCGGAAACCCAA
TCTTCCTCTATTCACGACTCGACCCTGTTCCCGAATCTCCCCGTTGGCAACATAAGCCCT
GTTTCCGAGAAGCAAGAAACGACCCCCTCTGCTTCTGATACAGAGACGATGAAGATGTTT
AATAATTGCGAAGAGGAGATGAATCGTGGAGAGACTAACTATGCTTATGAGAAACAGCAA
GAGGAGGAAGAAGAAGAAGATTGTGGAGGAGGAATGAGGATAGAGAGATCTGGAGATGGA
TATGTAATTAGATTGAAGTGTAGATGCAGACAAGCTTTTAGGGTTCTTTTCTCTGATCAT
CACCTCTACTTCAAGCCTCTC
>AtSKR Arabidopsis thaliana
ATGATAATGGAAGAAGAACTTCAAGAGCGTAACAACAATCTCAGCCTCGACGACATTGAT
CTTTCTCTCCTCCGTCTCACTTCACCACCGTACGACTACTCTCCTCCATCATCCTTGTTC
TCCGCCGATGAAATCACTCCTCCATTGAAACGCGCCTCCCCTGTTTCCGACGAATCCGAT
CTCCCCAAACGCAGAAAACTCTCCCCTCAAAACCCAATCTTTACCACTTCTCCTCCTCTC

TTCACTTCCCCGGAAACACAAACTTCCTCTGTTCATGACACGAGCCGGTACACGAATCTC
ACTAGCTCTGTTTCGGAGAAGCAAGAAGCGACGCCGTCTGTTACTGATACAGAGACGATG
AAGATGGTTAACAAATGCGTAGAGGTGATGAATCGTGGAGAGACTAACTATGCTTATGAG
AAGGAGCAAGAGGAGTTGGAAGAAGAAGAAGATTGTGGAGGAGGGATGAGGATAGAGAGA
TCTGGAGATGGGTTTGTAATTAGATTGAAGTGTAGATGCAGACAAGCTTTTAGGGTTCTT
TTCTCTGATGATCATCTCTACTTCAAGCCTCTC
>AlSKR Arabidopsis lyrata
ATGATGATGGAAGAACAACTTCAAGAGCGTAACCACAATCTGAGCCTCGAAGACATTGAT
CTATCTCTCCTCCGTCTCACTTCACCACGTGACTACTATTCTCCTTCATCATCCTTGTTC
TCCGCCGAGGAAATCACCCCTCCGTTGAAACGCGCCTCCCCTGTTTCCGACGAATCCGAT
CTCCCCAAACGCAGGAAAATCTCCCCTCAAAACCCAATCTTTGTCACCTCTCCTCGTCTC
TTCATTTCCCCGGAAACACAAACTTCCTCTGTTCATGACCCGACCCGGTACACGAATCTC
ACTAGCTCTGTTTCGGAAGAGCAAGAAACGACGCCGTCTGATCCTGATACAGAGACGATG
AAGATGGTTAACAATTGCGTTGAGAAGATGAAACGTGGAGAGACTAACTATGCTTATGAG
AAGGAGCAAGAGGAAGTGGAAGAAGAAGAAGAATGTGGAGGAGGGATGAGGATAGAGAGA
TCTGGAGATGGGTTTGTAATTAGATTGAAGTGTAAATGCAGACAAGCTTTTAGGGTTCTT
TTCTCTGATGATCATCTCTACTTTAAGCCTCTC
>EsSKR Eutrema salsugineum
ATGATGATGGAAGATGAATTTCAAGACCGTAACAACAATCTGAGCCTCGACGACATTGAT
CTCTCTCTCCTCCGCCTCTCTTCCCCACCTTGCGACTACTCTCGCTCCTCTTTCTTCTCC
GCCGTATCTCCTGAAATCAGCCCCTTGAAACGCTCCTCCCCTGTTTCCGACGAATCCGAT
CAGCCAAAACGCAGGAAGATTTCCCTCCAAAACCCAACCTTTATCACCTCTCCTCTTCGC
TTTACTTACCCGGAAACTCAATCTTCCTCAATCGACGACCAGACCCGGTACCCGAATCTC
ACCGCCGGCCAAGTCCGTTGCTCCCACGCAAGCCCTGTTTCCGAGAAGCACGAAACGTCG
CCGTCTGCTTCCAAGACTCCTGAGACAGAAACGATGAAGGTGGTTAACAATCGAGTAGAG
GAGATCAATCGCGAAGAGACTACTAACTCTGATGCTTACGAGAATCAGCAAAAGGAGGAA
GAAGATTGTGAAGAAGGGATGAGGATAGAGAGATCTGGAGATGGATTTGTAATTAGATTG
AAGTGTAGATGCAAACAAGCGTTTAGGGTTCTTTTCTCTGATCATCACCTCTACTTCAAG
CCTCTC
>CrSKR Capsella rubella
ATGATGATGATGATGATGGAAGAAAAAGCTGAAGATCGTAACCACAACCTGACCCTCGAT
GACATTGATCTCTCTCTCCTCCGTCTCACTTCCCCACCTTACGACTACTCACCCTTCTTA
TCCGCCGATGAAATCACCCCTCCGTTGAAACGCGCCTCCCCTGTTTCCGACGATTCCGAT
CAATCCAAACGAAGAAAACTCTCCCCTAAAGACTCGATCTTTATCACCTCTCCTGTTCTC
TTCACTTCCCCGGAGAACCAATCTTCATCTGTCCATGACCCGACTCAGTACACGAACCTC
ACAAGCCCTGTTTCGGAGAAGCAAGATACGACGCCGTCTGCTTCTGATACAGAGACGATG
AAGAAGGTTAGCAATTGCGTAGAGGAGATGAATCATGGAGAGACTAACTATGCTTATGAG
AAGGAGCAAGAGGAAGTGGAAGAAGAAGAAGAAGAAGAAGAAGATTGTGGAGGAGG
G
ATGAGGATAGAGAGATCTGGAGATGGGTTTGTAATTAGACTCAAGTGTAGTTGCAGACAA
GCTTTTAGGGTTCTTTTCTCTGATCATCACCTCTACTTCAAGGCTCTC
>BrSKR Brassica rapa
ATGATGATGATGGTGATGGAAAACGAGTTTCAAGAACGTAGCAACCTCAACATCAACGAC
ATTGATCTCTCTCTCCTCCGCCTCTCCTCCCCACCACCTTACGCCTACTCTCGCTCCTCC

TTCCTCTCCGCCGCATCTCCTGATGCTATCATCAGCCCCTTGAAACGCTCCTCCCCTGTT
TCCGACGAGTCAGATCAGCCCAAACGCAGGAGAGTCTCCCTCCAAGACCCCATCTTTATC
GCGTCTCCTCTCAGCTCCACTTACCCGGAAACTCAACCTTCCTCCATCGCCGACCCACCT
GTTCCGGAGAAGCCAGAAACGTCGCCGTATGATTCCAAGAGTCTTGAAACAGAGAGGATG
AAGATGGTTAACAATCACGTGGAGGAGATCAATCACGAAGAGACTAACTATGATACTTAT
GAAGAGATCTACCGAGAAGAGACTAACTATGATGCTTATGAGAAGCAGCAAGAAGAGGAG
GAAAAAGAAGAAGAGTGTGGAGAAGGGATGAGGATAGAGAGATCTGGAGATGGGTTTGTA
ATTAGATTGAAGTGTAGATGCAAGCTGGCTTACAGGGTTCTCTTCTCTGATCATCACCTC
TACTTCAAATCTCTC
>BnSKR Brassica napus
ATGATGATGATGGTAATGGAAAACGAGTTTCAAGAACGTAGCAATCTGAACATCAACGAC
ATTGATCTCTCTCTCCTCCGCCTCTCCTCCCCACCACCTTACGACTACTCTCGCTCCTCC
TTCCTCTCCGCCGCTTCTCCTGATGATATCAGCCCCTTGAAACGCTCCTCCCCTGTTTCC
GACGAGTCAGATCAGCCCAAACGCAGGAGAGTCTCCCTCCAAGACCCGATCTTTATCACG
TCTCCTCTCAGCTCCATCGACGACCCGACCCGCAAAAGCCCTGTTCCCGAGAAGCAAGAA
ACGTCGCCGTTTGATTCCAAGAGTTTTGAAACAGAGAGGATGAAGATGGTTAACAATCAC
GTTGAGGAGATCACTCACGAAGAGACTAACTATGATACTTATGAAGAGATCTACCGAGAA
GAGACTAACTATGATGCTTATGAGACGCAGCAAGAAGAAGAGTGTGGAGAAGGGATGAGG
ATAGAGAGATCTGGAGATGGATTTGTAATTAGATTGAAGTGTAGATGCAAGCTAGCTTAC
AGGATTCTCTTCTCTGATCATCACCTCTACTTCAAGTCCCTC
>RsSKR Raphanus sativus
ATGATGATGCAAGAAGAGTTTAAAGACCGTGCGAATCTGAGCATCAACGACATTGATCTC
TCACTCCTCCGCCTCTCCTCCCCACCAACTTACGACTCCTTCTTCTTCTCCCCCGTTTCT
CCTCCTCCTGATATCAGCCCCTTGAAACGCTCCTCCCCTGTTTCCGACGAATCCGATCAT
CCCAAACGCAGGAGACTCTCCTCTCAAGACCCAATCTTCATCACCTCTCCTCTTCGCTCC
TCCACCTATCATGAACCCCATTCCTCCTCTACCATCGAAGACCCGACTACTCACTACCCG
AATCTCGCCGCTGGCCACATCCTTTGTTCCCACTTAAGCTCTGTTTCCGAGAAGCAAGAA
ACGACGCTGTCTGCTTCCAAGACTCCTGATAAAGAGACGATGAAGATGGTTAACACTCAC
GTGGAGGAGATCGATCAAGAAGAGACTACTAACTATGATGCTTATGAGGAGATCAATCGC
GAGGAGACTAACTATGATGCTTATGAGGAAGAAAAAGAAGAAGAAGAAGATTGTGGAGAA
GGGATGAGGATAGAGAGATCTGGAGATGGGTTTGTTATTAGGTTGAAGTGTAGATGCAAG
CTAGCTTATAGGGTTCTCTTCTCTGATCACCACCTCTACTTCAAGTCTCTC
>CsSKR Camelina sativa
ATGATGAGGGAAGACAAAGTTATAGACCGTAACCATAATCTGAGCGTCAACGACATTGAT
CTCTCTCTGCTCCGTATCACTTCTCCTCCACCTTACTACGACTACTCTCCTTCCCCATCA
TTGTTACCCGCCGATGAAATCACACCTCCGTTGAAACGCGCCTCCCCTGTTTCCGACGAC
TCCGATCAGTCCAAACGAAGAAAACTCTCCCCTCAAGAAGAACCAATCTTTATAACCTCT
CCTCTTCTATTCACTTCCCGGGAGAACAACCAATCTTCTTCCTCTGTCCATGACCCGAAT
CTCACAAGCCCTGTTTTGGAGAAGCAAGATACGACGCCGTCTGCTTCTGATACAGAGACG
ATGAACAAGGTTAACCATTGCGTAGAGGAGATGAATCGGGGAGAGGTTAACTATGCTTAT
GAGCAAGAAGAAGTGGAACAAGAAGAAGAAGAAGAATGTGGAGGAGGGATGAGGATAGAG
AGATCAGGAGATGGTTTCGTAATTAGATTGAAGTGTAGATGCAGACAAGCTTTTAGGGTT
CTTTTCTCAGATCATCACCTCTACTTCAAGCCTCTC
>ThSKR Tarenaya hassleriana

ATGAAGATGAATTCGATCCATGAAGTTCTCTTCGAAGCCCATGAATCCTCGTTATCTGAC
ATGACAGAGCAAGATGGTTTTCATATCAACCTTGACGAAATCGATCTCTCTCTCCTCCGC
CTCAGCTCCTCCCCCTACAGCCTACGTGAATTTCTCCACCACTCGCCCGTCTCCGCTGCT
GCCGTCGCCATGAAAAGTCCTTCTCCTGACTCCGATGTCTCCGACCAGTCCAAACGCAGA
AAGATTTTAGTTCAAAACCAAGATTTGATCGACCCTTGTGTTTTTTCCGGCTACCCAGAG
AATCCACCTTCCTCCGAACCGACCCAGAATAGAACTCGCATCGGCGACCCGAGTCAACTC
GCTCCGGTCGTTCTTCGACGGTCGTTGTTTGATGGATACTTGTGCCCTCTTGCTGAGAAG
CAAGAAACGACGTTTAAGTGCAAGCCCTTCCCTGAAAAGACGATGATGGAGGAGAGTCCC
GAGTCAAAGATGCTAACGATGATAAACGATCGCGTGAAGGAGATGAACGGGTTGTGCAAG
GACTTGGATTTTCGCAGTGAAGAACACAGCCACAAAGAACTCGAAGATGCCACAAAGGAC
TTGGAGGAAGGTTTCGAAGAAGAAGGAATGAGAATAGAGAGATCCGGAGATGGATACGTG
ATCCGATTGAAGTGCCAATGCCGAAATGCTTATAGGGTTCTTTTCTCTGATAGTCACTTC
TACTTCAAGCTTCTA
>ChSKR Cardamine hirsuta
ATGATGACGGAAGACGAACTTCAAGAACGTAACAACAATCTTAGCCTCGACGACATTGAT
CTCTCCCTCCTTCGCCTCACTTCCCCTCCTTACGATCACTCATCGTCTTTCTTTTCCGCC
GATGAACTCAGCCCTCCTTTGAAACGCTCCTCCCCTGTTTCCAACGAATCCGATCACCAC
AAACGCAGAAAACTCTCCCCTCAAAACCCAATCTTTATTACCTCTCCTCTTCTCTTCACT
TCCCCGGAAACTCACGACTCGACTCTGTTCCCGAATCTCCCCGCCGGCAACATAAACCCT
GTTTCCGAGAAGCAAGAGACGACGCCGTCTGCTTATGATACAGAGACGATGAAGATGATT
AACAATTGCGAAGAGGAGATGAATCGCGGAGAGACTAACTATGCTTATGAGACGCAGCAA
GAGGAGGAAGAAGAAGAAGATTGTGGAGGAATGAGGATAGAAAGATCTGGAGATGGATAT
GTAATTAGATTGAAGTGTAGATGCAGAAAAGCTTTTAGGGTTCTTTTCTCTGATCATCAC
CTCTACTTCAAGCCTCTC
